# *Klebsiella pneumoniae* remodels its Kdo_2_-lipid A in a TLR4-dependent manner to adapt to the macrophage intracellular environment

**DOI:** 10.1101/2025.06.20.660740

**Authors:** Toby Leigh Bartholomew, Joana Sa-Pessoa, Rebecca Lancaster, Nan Zhang, Karolina Wojtania-O’Farrell, Steven J Hancock, Grant Mills, Adrien Kissenpfening, Jose A. Bengoechea

## Abstract

Pathogen adaptations to the intracellular macrophage environment remain poorly understood. We performed a high-resolution structural analysis of the lipid A purified from intracellular *Klebsiella pneumoniae* (KP). In both mouse and human macrophages, KP produces hexa- and hepta-acylated lipid A species modified with palmitate and 2-hydroxylated. LpxL1, PagP, and LpxO enzymes govern the intracellular lipid A in a PhoPQ-dependent manner triggered by the acidic pH of the KP-containing vacuole (KCV). Absence of LpxO and PagP lipid A modifications impairs intracellular survival and heightens NF-κB and IRF3-mediated inflammation, though KCV maturation remains unaffected. Instead, these modifications fortify the bacterial membrane. Absence of TLR4-TRAM-TRIF signalling increases KCV pH, impairing the production of the intracellular lipid A. KP intracellular survival is reduced in *tlr4^-/-^* macrophages, highlighting the critical role of this signalling pathway in KP immune evasion and how the pathogen has evolved to rely on innate immune system cues for virulence.

## INTRODUCTION

Macrophages have been at the heart of immune research for over a century and are an integral component of innate immunity. Macrophages sense pathogens through pattern recognition receptors facilitating phagocytosis and the activation of host defences upon the release of cytokines and chemokines. Among others, the family of Toll-like receptors (TLRs) play a pivotal role in recognition of microbial patterns such as the lipopolysaccharide (LPS) by TLR4, or bacterial flagellin by TLR5^1^. Activation of TLRs launches innate immune defence mechanisms controlled by NF-κB, IRF and MAP kinase signalling cascades^1,2^. Once phagocytosed, the microbe-containing vacuole follows a maturation process that results in the fusion of lysosomes and the exposition to oxygen-dependent and independent antimicrobial agents to eliminate the pathogen. However, despite these antimicrobial mechanisms, several pathogens have evolved strategies to overcome macrophages including avoidance of phagocytosis, perturbation of the phagolysosome maturation, shift of the macrophage plasticity towards a less antimicrobial state, and induction of cell death^3^. Bacterial factors such as capsule polysaccharides, type III and IV secretion systems, and pore-forming toxins have been demonstrated as pivotal anti-macrophage strategies. Nonetheless, the adaptations of pathogens to the intracellular macrophage environment remain poorly understood and, only recently, has started to be investigated using RNA-seq-based approaches^4–6^.

*Klebsiella pneumoniae* (KP) is a key contributor to the global rise in antibiotic-resistant infections, with both hospital- and community-acquired pneumonia caused by this pathogen representing a growing public health concern. Early on it was discovered the crucial role of alveolar macrophages in host defence against this pathogen^7^. This aligns with our findings demonstrating that the transcriptional pattern of KP-infected alveolar macrophages in vivo relates to TLR-controlled host defence responses governed by NF-κB^8^. Nonetheless, KP can flourish in the lung, suggesting that the pathogen may have evolved to withstand macrophages. Indeed, we demonstrated that KP survives intracellularly in mouse and human macrophages in a specific compartment, the KP containing vacuole (KCV), that deviates from the phagolysosome canonical maturation^9^.

Furthermore, KP induces a singular anti-inflammatory polarisation state, termed M(Kp), dependent on the activation of STAT6^8^. Absence of STAT6 limits KP intracellular survival and facilitates the clearance of the pathogen in vivo^8^, demonstrating that manipulation of macrophage biology plays an integral part in KP infection biology. However, the adaptations of KP to the intracellular environment within the KCV remain elusive.

To start addressing this gap in knowledge, we focus on determining the structure of the LPS lipid A section from intracellular KP. Gram-negative bacteria can modify its lipid A in response to environmental stimuli to fortify the membrane integrity and, therefore, it is tempting to hypothesize that KP may remodel its lipid A within the KCV as part of its adaptation to this harsh environment. We previously detailed KP lipid A structure produced by bacteria grown in vitro^10–12^. The main structure is a hexa-acylated lipid A corresponding to two glucosamines, two phosphates, four R-3-hydroxymyristoyl primary acyl chains, and two myristates (C_14_) as secondary acyl chains^10–12^ (Fig S1A). This lipid A species can be modified with the addition of palmitate to the hexa-acylated species (Fig S1B), by the hydroxylation of the C_14_ on the primary 2′-linked R-3-hydroxymyristoyl group (Fig S1C), and with the addition of 4CaminoC4CdeoxyCLCarabinose (L-Ara4N) and phosphoethanolamine (pEtN) to the 4′ and 1-phosphate, respectively (Fig S1D and Fig S1E). Interestingly, in the lungs of infected mice extracellular KP expressed two characteristic 2-hydroxylated hexa-acyl lipid A species that are lost in bacteria following in vitro minimally passage^13^. These in vivo lipid A modifications contribute to survival in vivo and mediate resistance to antimicrobial peptides^13^, providing support to the notion that KP remodels its lipid A to circumvent the innate immune system.

Here, we establish a method, we termed Intracell_lipid A_ method, to extract the lipid A from intracellular bacteria and analyse its structure by matrixCassisted laser desorption–ionisation timeCofCflight (MALDICTOF) mass spectrometry. We present evidence demonstrating that KP remodels its lipid A within the KCV upon activation of the two-component system PhoPQ, and that these lipid A modifications are necessary for intracellular survival. Furthermore, we revealed that TLR4-controlled modification of the KCV pH is the signal sensed by KP to remodel its lipid A. Altogether, this work illustrates how pathogens integrate cues from the innate immune system to promote virulence.

## RESULTS

### Analysis of *K. pneumoniae* intracellular lipid A

Previous studies have used the Bligh-Dyer method for lipid A extraction from intracellular bacteria^14,15^; however, when we used this method of purification in non-infected immortalized mouse bone-marrow-derived macrophages (iBMDMs) MALDI-TOF analysis revealed an ion of mass-to-charge (*m/z*) 1,963 (Fig S2A). Similarly, the same ion was detected using the Tri-Reagent method, another extraction protocol employed for lipid A analysis^16^ (Fig S2B). This host lipid has also been found by others in macrophages^15,17^. The fact that this host lipid could be interpreted as a KP hexa-acylated lipid A containing two glucosamines, two phosphates, four *R*-3-hydroxymyristoyl primary acyl chains, one myristate (C_14_), and one 2-hydroxymyristate (C_14:OH_) modified with phosphoethanolamine led us to develop a purification method by which this lipid is not detected while not affecting the isolation of KP lipid A. The new method, termed Intracell_lipid A,_ involves the lysis of cells with saponin before extraction of the lipid A from intracellular bacteria with the Tri-Reagent method. The protocol includes a detergent clean-up step to prevent contamination of the MALDI-TOF mass spectrometer. Using this method, ion *m/z* 1,963 was no longer detected in non-infected iBMDMs (Fig S2C). We next assessed whether the lipid A extract from KP grown in vitro using this method would be identical to those previously described^10–12^. We extracted and analysed the lipid A from strain CIP52.145 (hereafter Kp52145) grown in lysogeny broth (LB). This strain belongs to the KpI group and it encodes all virulence functions associated with invasive community-acquired disease in humans^18,19^. The lipid A contained the hexa-acylated species *m/z* 1,824 corresponding to two glucosamines, two phosphates, four 3COHCC_14_ and two C_14_, as well as two other peaks including *m/z* 1,840, corresponding to two glucosamines, two phosphates, four 3COHCC_14_, one C_14_ and one C_14:OH_, *m/z* 1,955 and *m/z* 2,063 corresponding to the addition of L-Ara4N (*m/z* 131) and palmitate (*m/z* 239), respectively, to the hexaCacylated (*m/z* 1,824) (Fig S2D), demonstrating that the Intracell_lipidA_ method extracts the same lipid A species as those reported before for KP grown in vitro.

We next applied this method to extract and analyse the lipid A from iBMDMs infected with Kp52145. Lipid A isolated from intracellular bacteria contained the hexa-acylated species *m/z* 1,840 and *m/z* 1,824 and the hepta-acylated species *m/z* 2,063, consistent with the addition of palmitate to ion *m/z* 1,824, found in vitro (Fig 1A). New lipid A species produced by intracellular bacteria were hexa-acylated species *m/z* 1,797, corresponding to two glucosamines, two phosphates, four 3COHCC_14_, one C_14_ and one laurate (C_12_), and *m/z* 1,813 consistent with four 3COHCC_14_, one C_12:OH_ and one C_14_ (Fig 1A). Other ions included *m/z* 2,036, *m/z* 2,052, and *m/z* 2,079, which are consistent with the addition of palmitate to ions *m/z* 1,797, *m/z* 1,813, and *m/z* 1,840, respectively (Fig 1A). Notably, the lipid A of bacteria recovered from the macrophages and grown on LB agar plates at 37^0^C for 24 h was identical to that from Kp52145 grown in LB (Fig 1B and Fig S2D), highlighting that the environment within the KCV dictates the lipid A produced by KP, which is lost in vitro. To demonstrate that the intracellular lipid A is not strain dependent, we analysed the lipid A produced by strain KP35. This strain clusters within the epidemic clonal group ST258 producing the KP carbapenemase^20^. In macrophages infected with KP35 we detected similar lipid A species as those found in Kp52145-infected cells (Fig S2E). The proposed lipid A structures are shown in Figure S3.

**Figure 1.**
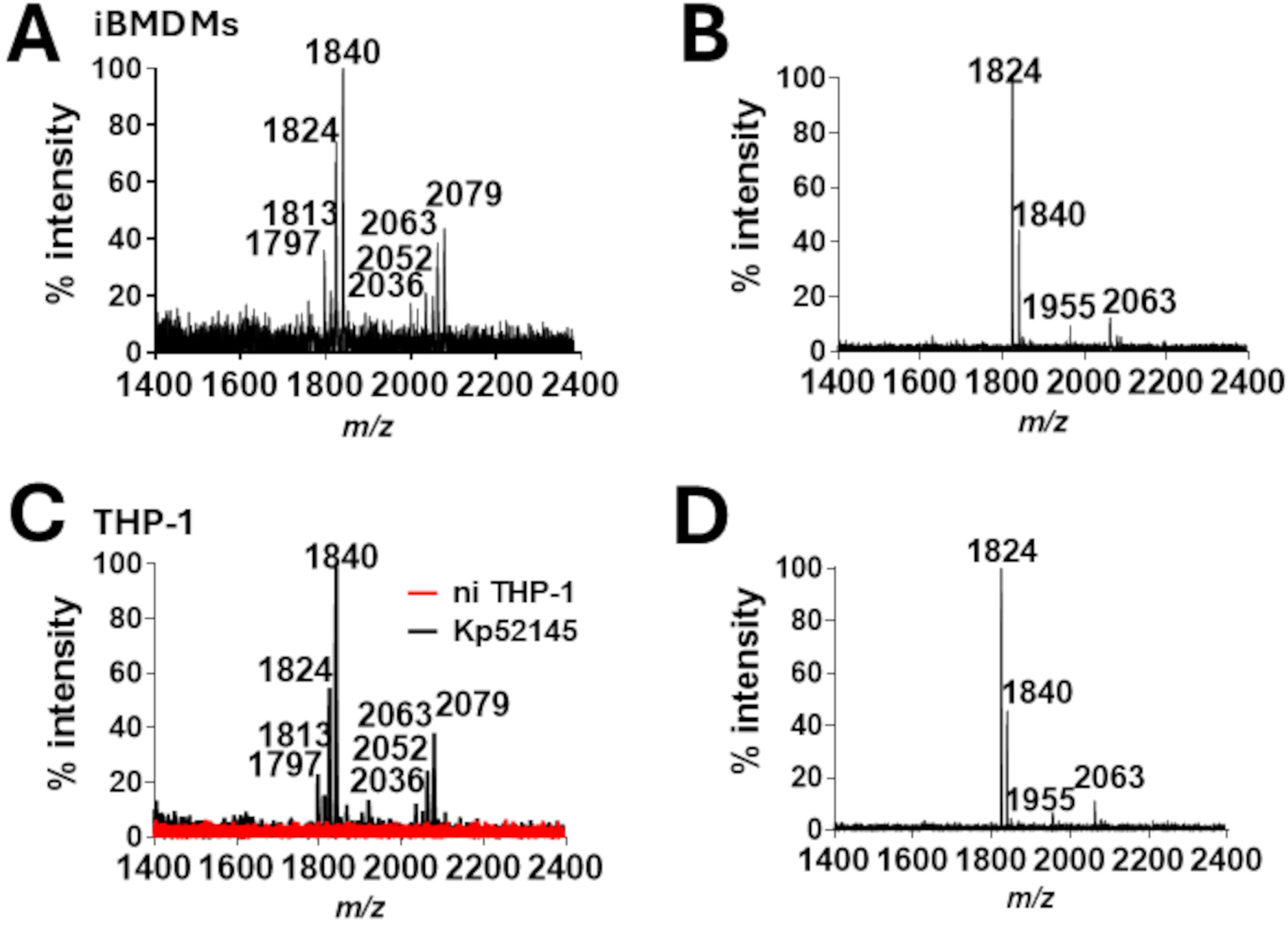
Lipid A produced by intracellular KP. Shown are negative-ion MALDI-TOF mass spectrometry spectra of lipid A purified from: A. intracellular Kp52145 in iBMDMs, B. Kp52145 grown for 24 h at 37°C in LB agar after being isolated from iBMDMs. C. intracellular Kp52145 in THP-1. D. Kp52145 grown for 24 h at 37°C in LB agar after being isolated from THP-1. Data represent the mass-to-charge ratios (*m/z*) of each lipid A species detected and are representative of three extractions.

We next sought to ascertain the lipid A pattern produced by intracellular KP in human macrophages. We extracted the lipid A from infected PMACdifferentiated THPC1 human macrophages. This cell line was derived from a patient with acute monocytic leukaemia and it is commonly used in human macrophage studies. The lipid A pattern produced by Kp52145 in PMA-treated THP-1 macrophages was identical to that found in infected iBMDMs (Fig 1C), demonstrating that the intracellular lipid A is not macrophage species-specific and suggesting that KP faces comparable intracellular environment in mouse and human macrophages. Likewise in the case of mouse macrophages, the lipid A from bacteria passaged once on LB agar plates was like that from KP grown in LB (Fig 1D and Fig S2D).

Altogether, this evidence demonstrates that KP extensively remodels its lipid A inside the KCV in both mouse and human macrophages. This intracellular lipid A contains new hexa-acylated species not found in vitro. The lipid A species are heavily modified with palmitate and 2-hydroxylated.

### Enzymology governing *K. pneumoniae* intracellular lipid A

To identify the enzymes responsible for the new lipid A species produced by intracellular KP we adopted a genetic approach. The hexa-acylated lipid A structure consists of a β(1′-6)-linked disaccharide of glucosamine phosphorylated at the 1 and 4′ positions, with positions 2, 3, 2′, and 3′ being acylated with R-3-hydroxymyristoyl groups, the so-called lipid IVA^21^. The 2′ and 3′ R-3-hydroxymyristoyl groups are further acylated^21^. Five enzymes are required to assemble the β(1′-6)-linked disaccharide that is characteristic of lipid A molecules^21^, whereas LpxK, KdtA, LpxL, and LpxM catalyse the last four enzymatic steps required to assemble the Kdo2–hexa-acylated lipid A^21^. LpxK phosphorylates the 4′ position of the disaccharide 1-phosphate to form lipid IVa; the next two Kdo residues are incorporated by the enzyme KdtA to generate the molecule Kdo2-lipid IVA^21^. The last steps involve the addition of the secondary fatty acid residues to the distal glucosamine unit by LpxL and LpxM, respectively, which require the Kdo disaccharide moiety in their substrates for activity^21^. Previous work demonstrated that KP LpxM catalyzes the transfer of C_14_ to the 3′ R-3-hydroxymyristoyl group^13^ and that LpxL2 is the acyltransferase that acylates the 2′ R-3-hydroxymyristoyl group with C_14_ to produce species *m/z* 1,824^12^. Therefore, we set out to identify the acyl transferase responsible for the species of the intracellular lipid A containing C_12_ at the 2′ R-3-hydroxymyristoyl group. Interestingly, KP encodes a second enzyme, LpxL1, that transfers C_12_ when expressed in the background of *Escherichia coli*^12^. To assess whether LpxL1 catalyses the acylation of the intracellular lipid A, iBMDMs were infected with the *lpxL1* mutant and the lipid A was purified and analysed as previously described. The lipid A produced by the *lpxL1* mutant lacked the species predicted to contain C_12_, *m/z* 1,797, *m/z* 1,813, *m/z* 2,036 and *m/z* 2,052 (Fig 2A).

**Figure 2.**
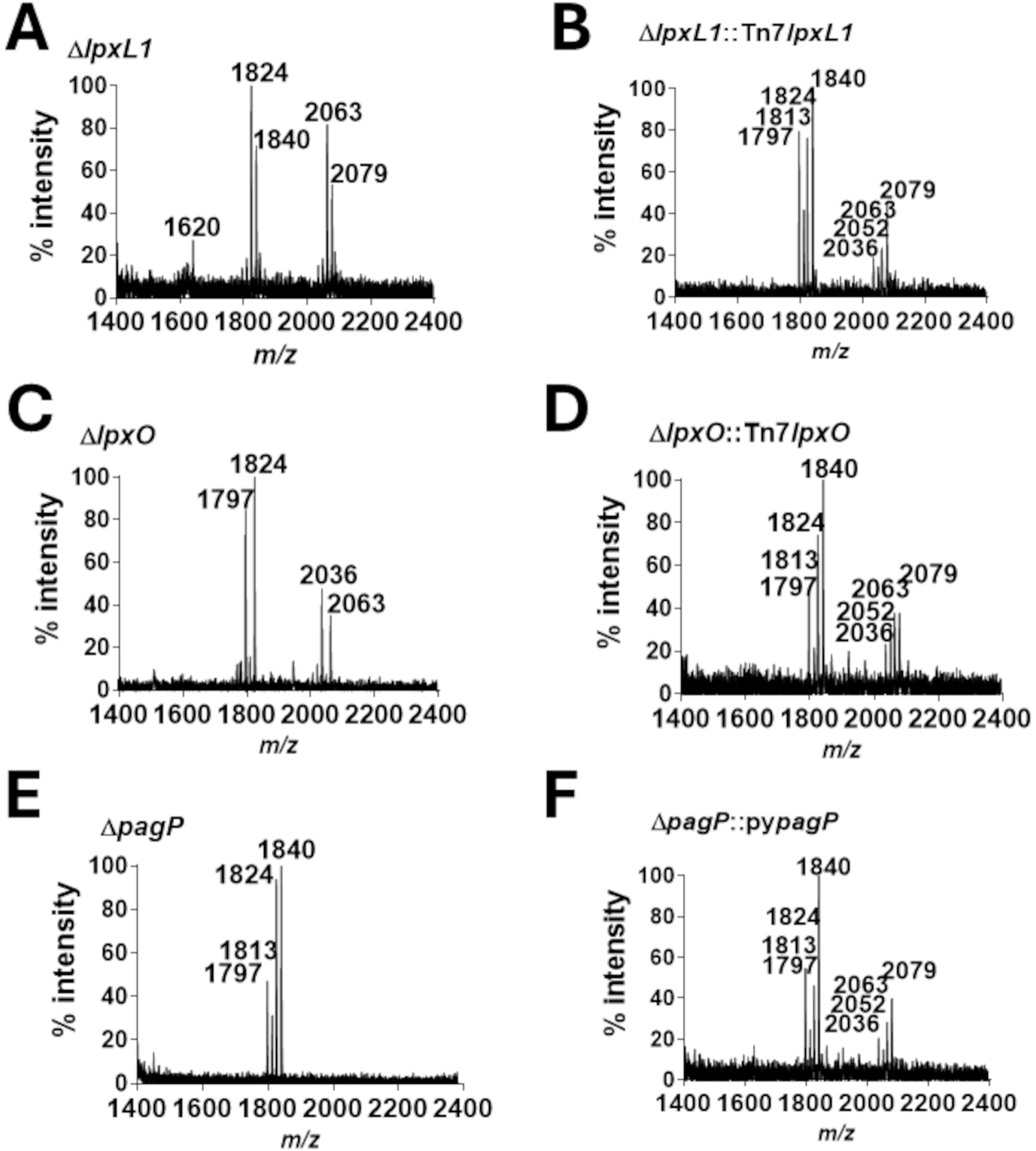
LpxL1, LpxO and PagP govern the lipid A produced by intracellular KP. Negative-ion MALDI-TOF mass spectrometry spectra of lipid A purified from intracellular bacteria: A. *lpxL1* mutant. B. Complemented *lpxL1* mutant C. *lpxO* mutant. D. Complementd *lpxO* mutant. E. *pagP* mutant. F. Complemented *pagP* mutant. Data represent the mass-to-charge ratios (*m/z*) of each lipid A species detected and are representative of three extractions.

Additionally, and as expected, a new species was detected, *m/z* 1,620, consistent with a penta-acylated lipid A corresponding to two glucosamines, two phosphates, four 3COHCC_14_, one C_14_ (Fig 2A). Complementation with LpxL1 restored the lipid A pattern (Fig 2B), demonstrating that LpxL1 is the acyltransferase responsible for the transfer of C_12_ to the to the acyl chain linked at the 2′ position of KP intracellular lipid A.

Another hallmark of the intracellular lipid A is the presence of hydroxylated species. We previously identified LpxO as the oxygenase that 2-hydroxylates KP lipid A^13^. We then ascertained whether LpxO mediates the hydroxylation of the intracellular lipid A. Analysis of the intracellular lipid A produced by the *lpxO* mutant revealed the absence of the predicted hydroxylated species *m/z* 1,813, *m/z* 1,848, *m/z* 2,052 and *m/z* 2,079 (Fig 2C). Complementation with LpxO restored the presence of the hydroxylated lipid A species (Fig 2D), confirming the crucial role of LpxO to govern the intracellular lipid A.

The fact that the species *m/z* 1,813 was not detected in the intracellular lipid A produced by *lpxl1* and *lpxO* mutants indicates that LpxO hydroxylates the C_12_ on the primary 2′-linked *R*-3-hydroxymyristoyl group. However, published evidence indicates that KP LpxO hydroxylates C_14_ on the same position^13^ (Fig 2C). These data suggest the intriguing notion that KP LpxO displays relaxed specificity for the type of fatty acid to hydroxylate and it has only specificity for the position of the acyl chain, the 2′ R-3-hydroxymyristoyl group. To provide evidence that KP LpxO hydroxylates the C_12_ at the 2′ R-3-hydroxymyristoyl group in addition to the C_14_ at this position, we followed a synthetic biology approach using *E. coli* as chassis to build *K. pneumoniae* lipid A. We first constructed a *lpxL* mutant in *E. coli* strain BN1 that produces hexa-acylated, bis-phosphorylated lipid A^12,22^ (Fig S4A and Fig S4B). We next expressed KP *lpxO* in the BN1 *lpxL* mutant background and, as anticipated, we did not observe any differences in the lipid A produced by this strain and the *lpxL* mutant (Fig S4C). This finding is consistent with our previous evidence demonstrating that KP LpxO does not hydroxylate the acyl chain transferred by LpxM on the primary 3′ R-3-hydroxymyristoyl group^13^. When we complemented the *E. coli lpxL* mutant strains with KP *lpxL1*, we observed the presence of species *m/z* 1,813 in the strain expressing *lpxO* (Fig S4D), confirming that LpxO can hydroxylate the C_12_ at the 2′ R-3-hydroxymyristoyl group, and lipid A species *m/z* 1,797 corresponding to two glucosamines, two phosphates, four 3COHCC_14_, one C_14_ and one C_12_ acylating the 2′ R-3-hydroxymyristoyl group (Fig S4D and Fig S4E).

Lastly, we set out to resolve the lipid A species of the hepta-acylated species of the intracellular lipid A predicted to contain palmitate. PagP is the acyltransferase responsible for adding palmitate to KP lipid A^10,11^. We then infected iBMDMs with a *pagP* mutant and determined the lipid A produced by this mutant. As anticipated, the intracellular lipid A lacked the predicted palmitoylated lipid A species *m/z* 2,036, *m/z* 2,052, *m/z* 2,063 and *m/z* 2,079 (Fig 2E). Complementation with *pagP* restored the presence of palmitoylated species (Fig 2F), corroborating that PagP is the acyltransferase responsible for the hepta-acylated species of the intracellular lipid A containing palmitate.

PhoPQ controls *K. pneumoniae* intracellular lipid A.

The two-component systems PhoPQ and PmrAB control the lipid A produced by KP in vitro^10^. Therefore, we asked whether these two-components systems also govern the intracellular lipid A. iBMDMs were infected with the *phoQ* and *pmrAB* mutants, and the lipid A was extracted and analysed by MALDI-TOF. The lipid A produced by the *pmrAB* mutant was like that produced by the wild-type strain (Fig 3A) and contained the previously described hexa-acylated and hepta-acylated lipid A species . In contrast, the lipid A produced by the *phoQ* mutant lacked species *m/z* 1,813, *m/z* 2,036, *m/z* 2,052, *m/z* 2,063 and *m/z* 2,079 (Fig 3B). Of note, this lipid A resembles that produced by the wild-type strain grown in LB (Fig S2D), suggesting that absence of PhoPQ impedes KP to remodel its lipid A in the KCV. Complementation restored the intracellular lipid A pattern found in the wild-type strain (Fig 3C).

**Figure 3.**
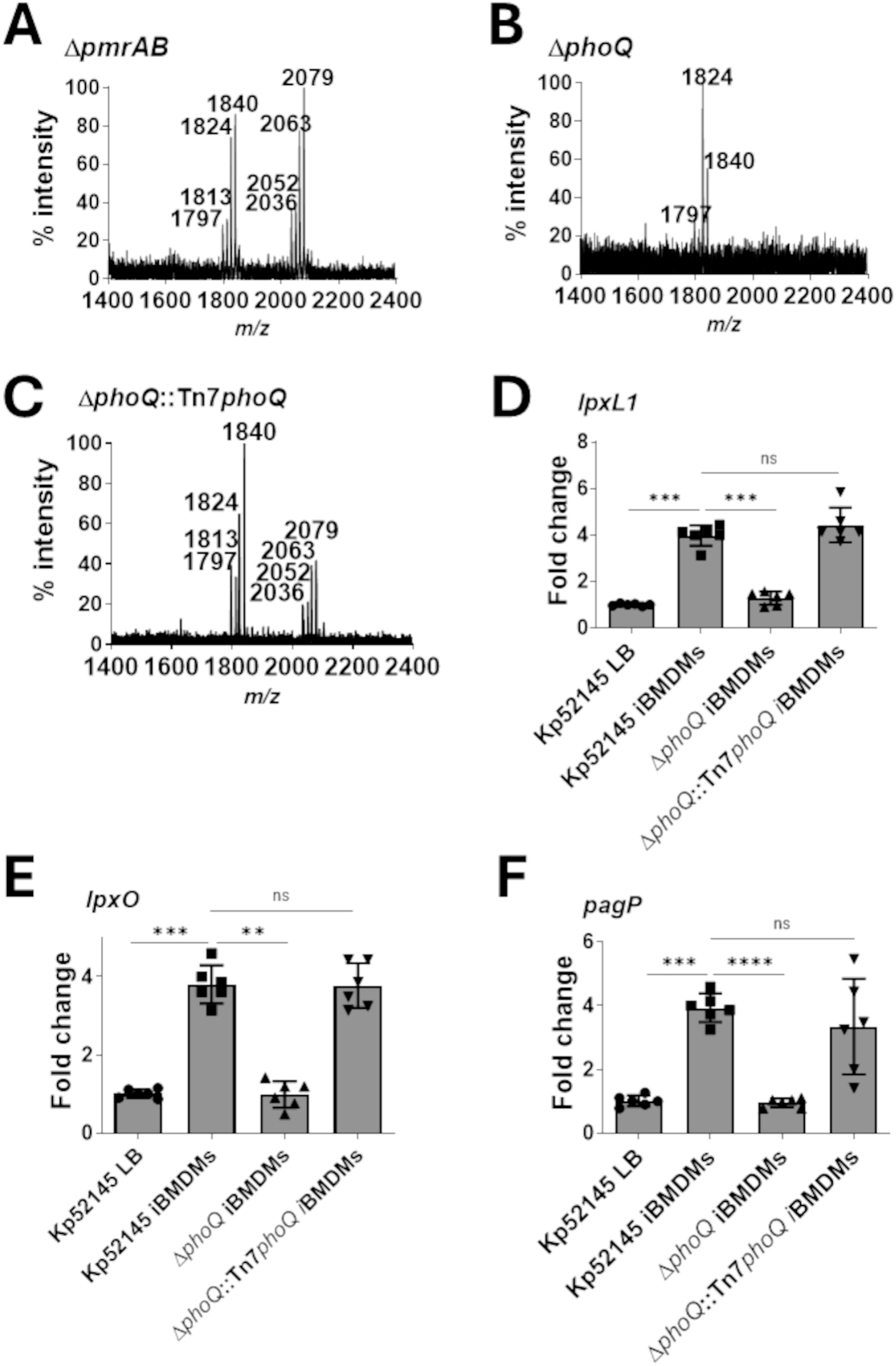
PhoPQ controls the lipid A produced by intracellular KP. A. Negative-ion MALDI-TOF mass spectrometry spectra of lipid A purified from intracellular *pmrAB* mutant. B. Negative-ion MALDI-TOF mass spectrometry spectra of lipid A purified from intracellular *phoQ* mutant. C. Negative-ion MALDI-TOF mass spectrometry spectra of lipid A purified from intracellular complemented *phoQ* mutant. D. *lpxL1* expression by wild-type KP and the *phoQ* mutant KP grown in LB or intracellularly in iBMDMs determined by reverse transcriptase quantitative realCtime PCR (RT-qPCR). E. *lpxO* expression by wild-type KP and the *phoQ* mutant KP grown in LB or intracellularly in iBMDMs determined by RT-qPCR. F. *pagP* expression by wild-type KP and the *phoQ* mutant KP grown in LB or intracellularly in iBMDMs determined by RT-qPCR. In panels A, B and C data represent the mass-to-charge ratios (*m/z*) of each lipid A species detected and are representative of three extractions. In panels D, E, and F values are presented as the means ± SD from three independent cDNA preparations measured in duplicate. *P* values were ns (not significant), <0.01 (**), <0.001 (***) and <0.0001 (****) for the indicated comparisons determined using one-way ANOVA with Bonferroni contrasts.

We next investigated whether PhoPQ controls the expression of the enzymes responsible for the intracellular lipid A. iBMDMs were infected with the wild-type strain, the *phoQ* and the complemented strain and bacterial mRNA was isolated to determine the expressions of *lpxL1*, *lpxO* and *pagP*. As reference, we extracted the mRNA from Kp52145 grown in LB to mid-exponential phase. This type of sample is used in studies as a comparator to determine the transcriptome of intracellular bacteria^4^. RT-qPCR experiments showed that the expressions of *lpxL1*, *lpxO* and *pagP* were higher in intracellular bacteria than in those grown in LB (Fig 3D). PhoPQ controlled the expressions of these genes in intracellular bacteria because their expressions decrease in the *phoQ* mutant to the levels found in bacteria grown in vitro (Fig 3D). Complementation restored the expressions of *lpxL1*, *lpxO* and *pagP* to the levels found in Kp52145-infected macrophages (Fig 3D).

Altogether, this evidence establishes that KP remodels its lipid A inside the KCV in a PhoPQ-dependent manner.

### LpxO and PagP modifications are required for intracellular survival

We next sought to determine the contribution of the lipid A produced inside the KCV to intracellular survival. We infected iBMDMs with *lpxO* and *pagP* mutants to assess whether the absence of LpxO and PagP-dependent lipid A modifications impairs KP intracellular survival. We did not observe any differences between any of the strains in the attachment to macrophages (Fig 4A) whereas the *lpxO* and *pagP* mutant were engulfed in significantly higher numbers than the wild-type strain (Fig 4B). Time-course experiments revealed that the intracellular survival of *lpxO* and *pagP* mutants were significantly reduced (Fig 4C). Complementation of the *lpxO* and *pagP* mutants restored the phagocytosis and intracellular survival to wild-type levels (Fig 4B and Fig 4C).

**Figure 4.**
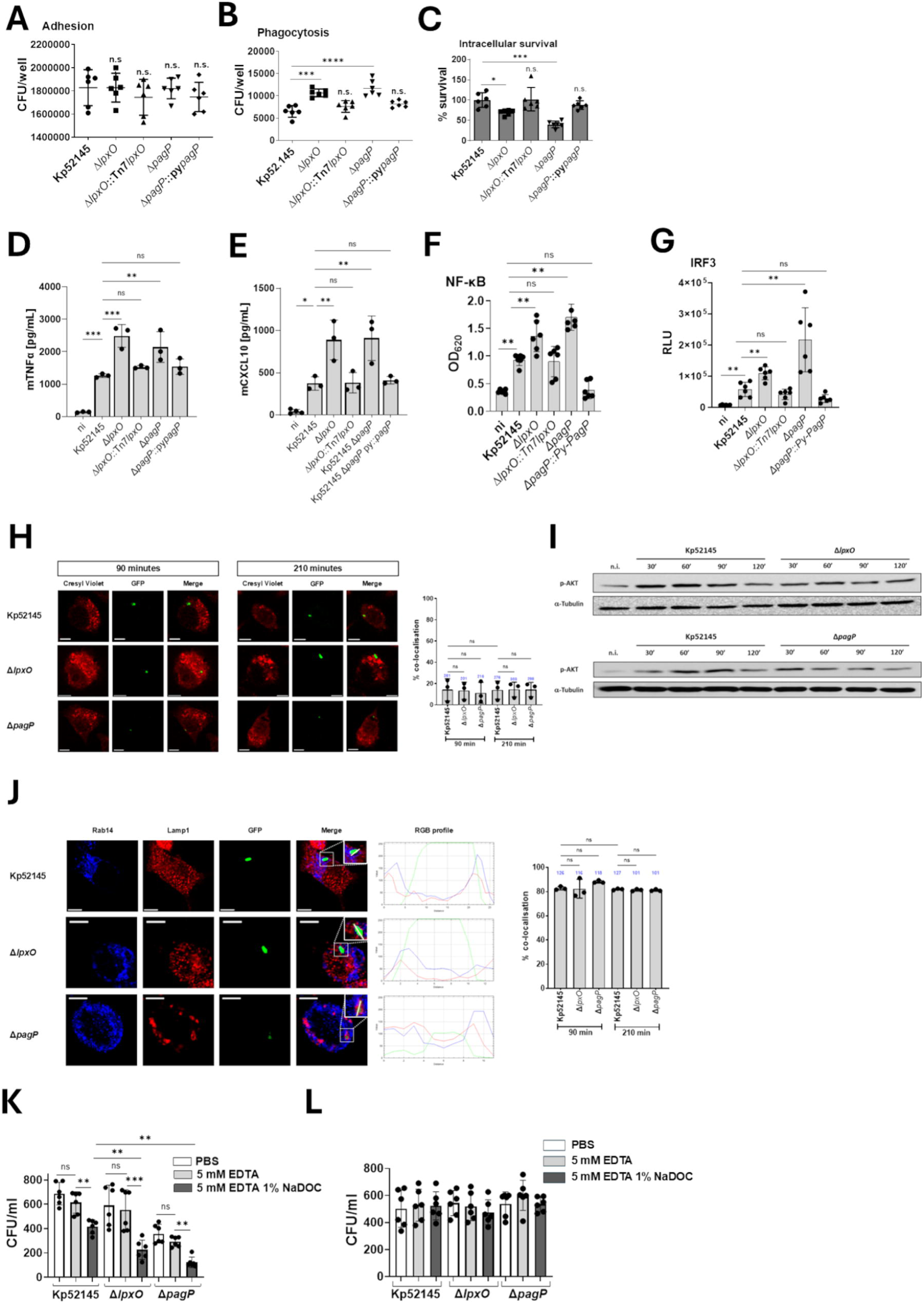
Role of LpxO and PagP-dependent lipid A modifications on KP intracellular survival. A. Adhesion of Kp52145, *lpxO* and *pagP* mutants, and their complemented strains to wild-type iBMDMs. B. Phagocytosis of Kp52145, *lpxO* and *pagP* mutants, and their complemented strains by wild-type iBMDMs. C. Kp52145, *lpxO* and *pagP* mutants, and their complemented strains intracellular survival in wild-type iBMDMs 5 h after addition of gentamycin. Results are expressed as % of survival (CFUs at 5.5 h versus 1.5 h normalized to the results obtained in wild-type macrophages infected with Kp52145 set to 100%). D. Secretion of TNFα by infected wild-type iBMDMs with Kp52145, *lpxO* and *pagP* mutants, and their complemented strains. E. Secretion of CXCL10 by infected wild-type iBMDMs with Kp52145, *lpxO* and *pagP* mutants, and their complemented strains. F. Activation of the NF-κB signalling pathway in infected THP-1 Dual macrophages with Kp52145, *lpxO* and *pagP* mutants and their complemented strains measured by assessing the levels of secreted SEAP reporter enzyme. G. Activation of the IRF3 signalling pathway in infected THP-1 Dual macrophages with Kp52145, *lpxO* and *pagP* mutants and their complemented strains measured by assessing the levels of secreted luciferase Lucia luciferase reporter enzyme. H. Immunofluorescence confocal microscopy of the coClocalisation of KP strains harbouring pFPV25.1Cm, and cresyl violet dye in wildCtype iBMDMs. The images were taken 90Cmin and 210 min post infection. Images are representative of duplicate coverslips of three independent experiments. Panel include the percentage of KCV coClocalisation with cresyl violet over a time course. The number of KCVs assessed per strain and time point are shown on top of each bar. I. Immunoblot analysis of phosphorylated AKT (p-AKT) and tubulin levels in lysates from nonCinfected (ni) and infected wild-type iBMDMs with Kp52145, *lpxO* and *pagP* mutants. J. Immunofluorescence confocal microscopy of the coClocalisation of the KCV with Rab14, and Lamp1, KCV marker^9^, in infected wild-type iBMDMs with KP strains harbouring pFPV25.1. The images were taken 210 min post infection. Images are representative of duplicate coverslips of three independent experiments. Panel includes the RGB profile (RGB Profiler-ImageJ plugging) showing the intensity changes along the length of the line. Panel include the percentage of KCV coClocalisation with Rab14 over a time course. The number of KCVs assessed per strain and time point are shown on top of each bar. K. Susceptibility of intracellular KP to sodium deoxycholate (NaDOC). Infected wild-type iBMDMs with Kp52145, *lpxO* and *pagP* mutants were lysed, and bacteria were incubated with PBS, EDTA, or EDTA-NaDOC for 1 h. Bacteria were quantified after serial dilution followed by plating on LB agar plates. L. Susceptibility of KP grown in LB to NaDOC. Exponentially grown KP strains in LB were incubated with PBS, EDTA, or EDTA-NaDOC for 1 h. Bacteria were quantified after serial dilution followed by plating on LB agar plates. Error bars are presented as the meanC±CSD of three independent experiments in duplicate. Immunoblot images are representative of three independent experiments. Microscopy images are representative of duplicate coverslips of three independent experiments. *P* values were ns (not significant), <0.05 (*), <0.01 (**), <0.001 (***) and <0.0001 (****) for the indicated comparisons determined using one-way ANOVA with Bonferroni contrasts.

This reduced intracellular survival was associated with an increase in the inflammatory response because cells infected with the *lpxO* and *pagP* mutants secreted higher levels of the MyeloidCDifferentiation factorC88 (MyD88)-dependent cytokine tumor necrosis factor α (TNF-α) and of the Toll/ILC1R domainCcontaining adaptorCinducing IFNCβ (TRIF)-dependent cytokine CXCL10 than macrophages infected with the wild-type strain (Fig 4D and Fig 4E). Complementation of the mutants restored the secretion of cytokines by infected cells with the mutants to the levels secreted by cells infected with the wild-type strain (Fig 4D and Fig 4E), demonstrating that absence of the LpxO and PagP-dependent lipid A modifications results in heightened inflammatory response upon infection. We next sought to identify the signalling pathways governing this inflammatory response. We infected THP1-Dual macrophages that allow the assessment of the activation of the NF-κB pathway governing MyD88 responses, by monitoring the activity of an NF-κB-inducible SEAP reporter construct, and the IRF3 pathway controlling TRIF responses, by determining the levels of secreted luciferase (Lucia) reporter gene under the control of IFN-stimulated response elements (ISRE) fused to an ISG54 minimal promoter. This promoter is unresponsive to activators of the NF-kB or AP-1 pathways. NF-κB activation was significantly higher in cells infected with the *lpxO* and *pagP* mutants than in those infected with the wild-type strain (Fig 4F). Similar results were observed when the activation of IRF was investigated (Fig 4G). Complementation of the mutants restored the activation of NF-κB and IRF3 to the levesl induced by the wild-type strain (Fig 4F and Fig 4G), indicating that absence of LpxO and PagP triggers a heightened activation of the NF-κB and IRF3 signalling pathways controlling inflammation in macrophages.

To explain the reduced intracellular survival of the *lpxO* and *pagP* mutants, we hypothesized that the absence of LpxO and PagP results in an increase colocalization of lysosomes with the KCV. Lysosomes were labelled with the membrane-permeant fluorophore cresyl violet^23^, and cells were infected with GFP-labelled bacteria to assess the KCV at the single-cell level by immunofluorescence. Confocal microscopy experiments revealed that the majority of the KCVs from infected macrophages did not colocalize with cresyl violet (Fig 4H). KP targets the PI3K-AKT axis to survive intracellularly by triggering the recruitment of Rab14 to the KCV to block the fusion with lysosomes^9^. We found no differences between the *lpxO* and *pagP* mutants and the wild type in the activation of AKT phosphorylation (Fig 4I). We also observed no differences in the recruitment of Rab14 to the KCV in macrophages infected with the wild-type strain, and *lpxO* and *pagP* mutants (Fig 4J). Altogether, this evidence refuted the hypothesis that absence of LpxO and PagP perturbs the maturation of the KCV.

We then speculated that the lipid A modifications controlled by both enzymes may be needed to ensure KP viability within the KCV. The KCV is most likely a harsh environment due to the low pH and the limited availability of ions and nutrients. We then asked whether the fitness of the *lpxO* and *pagP* mutants is compromised in media with varying pH and ion content. We did not detect any growth differences between the wild type and the mutants in LB either at pH 7.0 or at pH 5.5 (Fig S5A and Fig S5B). In M9-0.2% glucose the *pagP* mutant showed a growth defect at pH 7.0, pH 5.5 and pH 4.8 (Fig S5C Fig S5D and Fig S5E) whereas the *lpxO* mutant only showed a growth defect when grown at pH 4.8 (Fig S5E). When we grew the strains in M9-0.2% glucose with low magnesium buffered to pH 5.0 (Fig S5F) and the PCN medium, reported to recapitulate signals found by Gram-negative pathogens in the pathogen containing vacuole^4,24^, with high levels K_2_HPO_4_/KH_2_PO_4_ buffered to pH 5.0 (Fig S5G), only the *pagP* mutant showed a growth defect. Altogether, these findings illustrate that PagP and LpxO-dependent modifications, particularly the former, are important for KP survival in acidic pH in nutrient restricted conditions.

The acidic pH and the low amount of ions are also known to negatively affect the bacterial membranes^25,26^. Therefore, we next considered whether the LpxO and PagP modifications contribute to fortify the membrane barrier within the KCV. To investigate this notion, macrophages were infected with the wild type and the *lpxO* and *pagP* mutants, and the recovered intracellular bacteria were exposed to the detergent sodium deoxycholate (NaDOC) in PBS buffer containing EDTA. NaDOC is excluded by the membrane of Gram-negative bacteria and an increased susceptibility is an indication of a weakened membrane barrier^27^. Exposure of intracellular Kp52145 to NaDOC resulted in a 30% decrease in bacteria levels compare with bacteria exposed to vehicle solution for the detergent (Fig 4K), suggesting that intracellular KP is susceptible to detergents. This is consistent with previous observations showing that for recovery of intracellular KP from infected macrophages mild detergents like saponin should be used instead of Triton X-100^9^. The intracellular *lpxO* and *pagP* mutant bacteria were more susceptible to NaDOC than the wild type (Fig 4K). 60 and 58% less *lpxO* and *pagP* less bacteria were recovered after NaDOC treatment, respectively, (Fig 4K). Incubation with EDTA had no effect on any of the strains (Fig 4K). Importantly, these findings were related to the intracellular environment encountered by the strains because neither Kp52145 nor the *lpxO* and *pagP* mutants were susceptible to NaDOC when grown in LB (Fig 4L). This evidence demonstrates that LpxO and PagP-controlled remodelling of the lipid A is necessary for KP survival within the KCV by fortifying the bacterial membrane barrier.

### The KCV acidic pH is the signal inducing the intracellular lipid A

A defining characteristic of the KCV is its acidic pH^9^, and KP intracellular survival requires the acidification of the KCV^9^. Therefore, we asked whether the pH of the vacuole could be a stimulus for inducing the intracellular lipid A. Treatment of macrophages with bafilomycin A1, a specific inhibitor of the vacuolar type H+-ATPase in cells, inhibits the acidification of organelles containing this enzyme, such as lysosomes and endosomes^28^. Corroborating previous observations^9^, control experiments showed that bafilomycin A1 abrogated the overlap between KP and the fixable acidotropic probe LysoTracker (Fig S6). We then treated the macrophages with bafilomycin A1 and extracted the lipid A from the intracellular bacteria. The lipid A produced by Kp52145 from bafilomycin A1-treated macrophages contained the hexa-acylated species *m/z* 1,797, *m/z* 1,813 and *m/z* 1.824 (Fig 5A). The only hydroxylated species detected was *m/z* 1,840 whereas no hepta-acylated species containing palmitate were noted (Fig 5A). Therefore, the acidification of the KCV is a cue exploited by *Klebsiella* to remodel the lipid A within the KCV.

**Figure 5.**
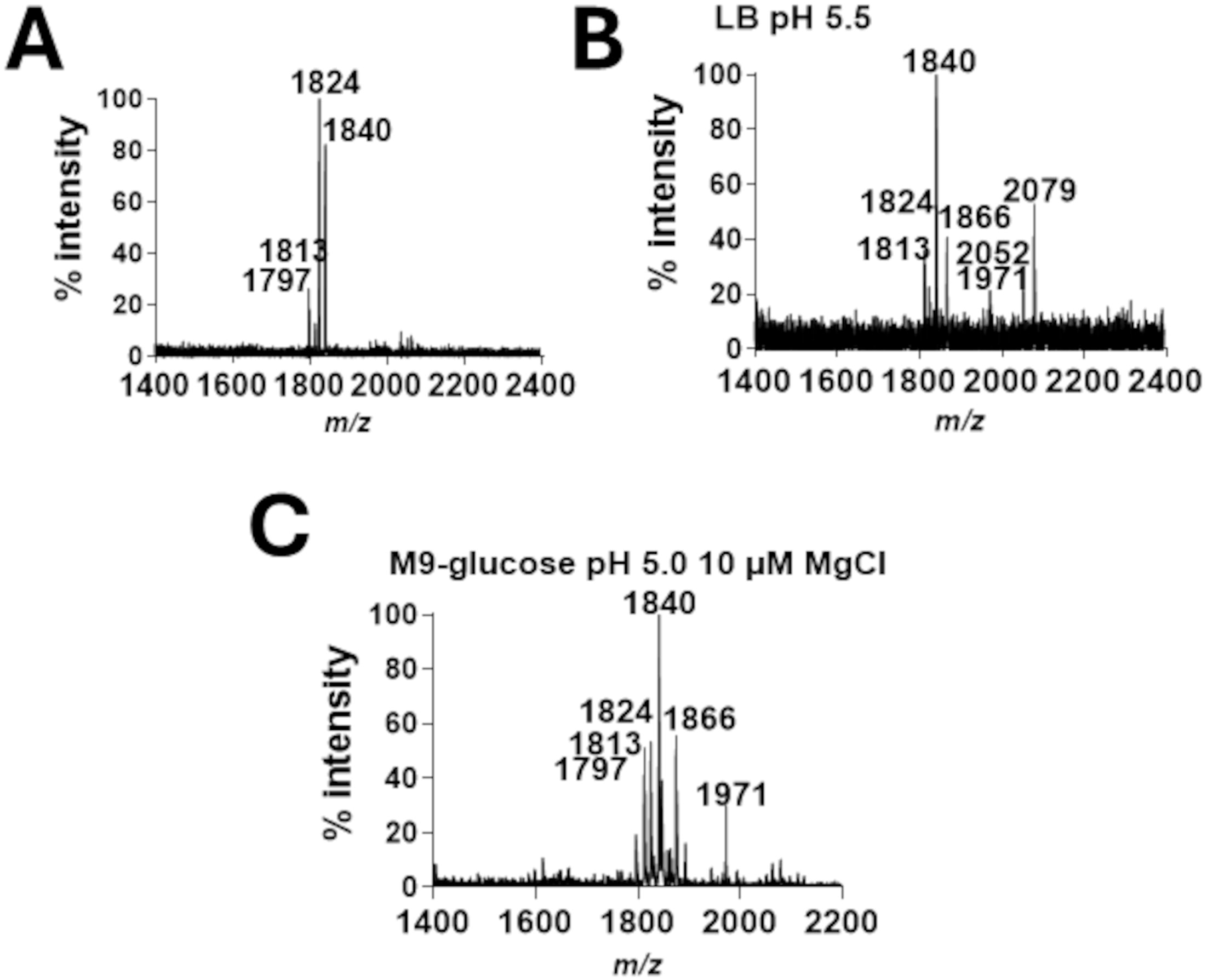
Effect of the KCV pH on the intracellular KP lipid A. Negative-ion MALDI-TOF mass spectrometry spectra of lipid A purified from: A. Intracellular Kp52145 in bafilomycin A1-treated iBMDMs. B. Kp52145 grown in LB buffered to pH 5.5 C. Kp52145 grown in M9-glucose buffered to pH 5 with low magnesium (10 µM MgCl_2_). In panels A, B and C data represent the mass-to-charge ratios (*m/z*) of each lipid A species detected and are representative of three extractions.

We next assessed whether an acidic pH is enough to trigger the intracellular lipid A pattern in vitro. Kp52145 grown in LB at pH 5.5 produced a lipid A containing species *m/z* 1,824 and the two hydroxylated species *m/z* 1,813 and *m/z* 1.840 and the hepta-acylated *m/z* 2,079 because of the addition of palmitate to species *m/z* 1,840 (Fig 5B). The hexa-acylated species *m/z* 1,797 and the hepta-acylated *m/z* 2,036, *m/z* 2,052 and *m/z* 2,063, that are found in the intracellular lipid A, were not detected. Instead, two new species were observed. Species *m/z* 1,866, a hexa-acylated lipid A previously described in KP^13^, and *m/z* 1,971 consistent with the addition of 4CaminoC4CdeoxyCLCarabinose (*m/z* 131) to *m/z* 1,840. The lipid A produced by KP grown in M9 glucose buffered to pH 5.0 and containing low magnesium also lacked the hepta-acylated species (Fig 5C). Altogether, these results indicate that the environment encountered by KP within the KCV cannot be fully recapitulated in vitro.

### TLR4 signalling controls the intracellular lipid A

The importance of TLR signalling in sensing KP infections has previously been shown^29^. We sought to test the effect of TLR deficiency on the production of the intracellular lipid A by KP. We first tested macrophages double knock-out for *tlr2* and *tlr4*. Both receptors have been demonstrated to control host defence responses against KP^30,31^. The lipid A produced by Kp52145 in these macrophages lacked the species *m/z* 1,813 and the hepta-acylated species *m/z* 2,036, *m/z* 2,052 and *m/z* 2,063 (Fig 6A). We next infected single knock-out cells for *tlr2* and *tlr4* to determine the relative role of each receptor. Interestingly, in *tlr2*^-/-^ macrophages, Kp52145 produced the characteristic intracellular lipid A (Fig 6B) whereas it was lost in infected *tlr4*^-/-^ macrophages (Fig 6C), demonstrating that TLR4 signalling governs the production of the intracellular lipid A.

**Figure 6.**
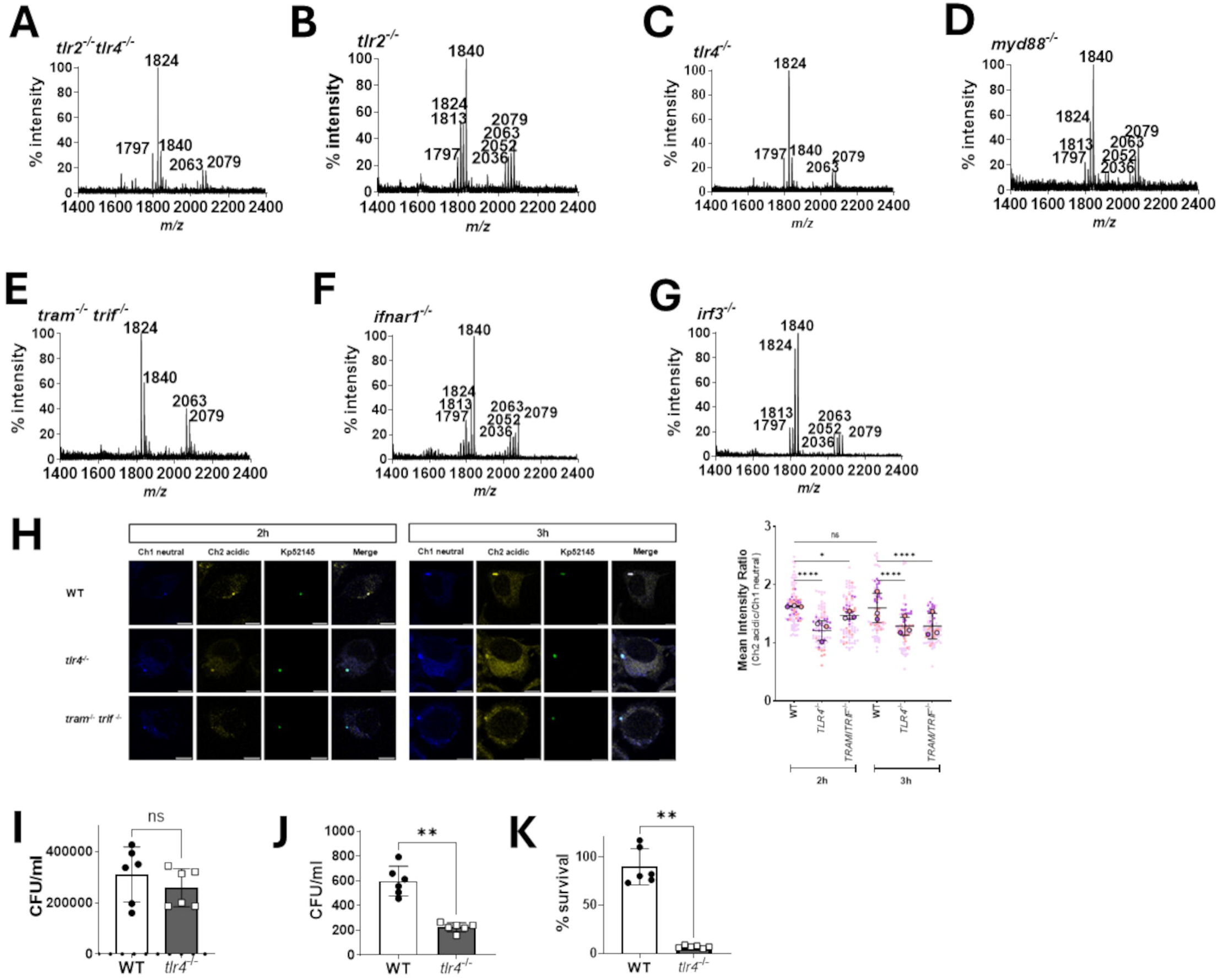
TLR4 signalling controls the intracellular lipid A. A. Negative-ion MALDI-TOF mass spectrometry spectra of lipid A purified from intracellular Kp52145 in *tlr2*^-/-^ -*tlr4*^-/-^ iBMDMs. B. Negative-ion MALDI-TOF mass spectrometry spectra of lipid A purified from intracellular Kp52145 in *tlr2*^--/-^ iBMDMs. C. Negative-ion MALDI-TOF mass spectrometry spectra of lipid A purified from intracellular Kp52145 in *tlr4*^--/-^ iBMDMs. D. Negative-ion MALDI-TOF mass spectrometry spectra of lipid A purified from intracellular Kp52145 in *myd88*^--/-^ iBMDMs. E. Negative-ion MALDI-TOF mass spectrometry spectra of lipid A purified from intracellular Kp52145 in *tram*^-/-^ -*trif*^-/-^ iBMDMs. F. Negative-ion MALDI-TOF mass spectrometry spectra of lipid A purified from intracellular Kp52145 in *ifnar1*^--/-^ iBMDMs. G. Negative-ion MALDI-TOF mass spectrometry spectra of lipid A purified from intracellular Kp52145 in *irf3*^--/-^ iBMDMs. H. Confocal microscopy-based assessment of the pH of the KCV using the radiometric probe LysoSenor. Wild-type, *tlr4*^--/-^ and *tram*^-/-^ -*trif*^-/-^ iBMDMs were infected with Kp52145 harbouring pFPV25.1Cm, and images were taken 2 and 3 h post infection. Images are representative of duplicate coverslips of five independent experiments. Panel includes the super plot analysis of the mean intensity ratio of the KCV. I. Adhesion of Kp52145, to wild-type and *tlr4*^--/-^ iBMDMs. J. Phagocytosis of Kp52145 by wild-type and *tlr4*^--/-^ iBMDMs. K. Kp52145 intracellular survival in wild-type and *tlr4*^--/-^ iBMDMs. 5 h after addition of gentamycin. Results are expressed as % of survival (CFUs at 5.5 h versus 1.5 h normalized to the results obtained in wild-type macrophages infected with Kp52145 set to 100%).

TLR signalling involves a series of different adaptors. MyD88 is a universal adaptor used by all TLRs except TLR3, TRIF is used by TLR3 and TLR4, whereas TRIFCrelated adaptor molecule (TRAM) is recruited by endosomal located TLR9 and TLR4. Therefore, we asked whether MyD88, TRAM and TRIF govern the production of the intracellular lipid A^32^. Infection of *myd88^-/-^* macrophages did not affect the production of the intracellular lipid A (Fig 6D). In contrast, the lipid A produced by Kp52145 in *tram-trif^-/-^* macrophages lacked the hexa-acylated species *m/z* 1,797 and *m/z* 1,813, and the hepta-acylated species *m/z* 2,036, and *m/z* 2,052 (Fig 6E), indicating that absence of TRAM-TRIF impairs the production of the intracellular lipid A.

The fact that TLR4-TRAM-TRIF controls the production of type I IFN upon KP infection^33^, led us to ascertain the effect of type I IFN signalling on the intracellular lipid A. We then infected type I IFN receptorCdeficient (*ifnar1^-/-^)* macrophages and purified and analysed the lipid A. Absence of IFNAR1 did not affect the production of the intracellular lipid A by Kp52145 (Fig 6F). Likewise, absence of IRF3, the factor controlling type I IFN production upon KP infection^33^, did not impair the ability of KP to produce the intracellular lipid A pattern (Fig 6G). Together, this evidence indicates that type I IFN signalling is not required for the expression of the intracellular lipid A.

We next tested whether the absence of other innate proteins that have been demonstrated crucial in KP infection biology, IL10, the absent in melanoma 2 (AIM2) inflammasome and its adaptor ASC, and the sterile α and HEAT armadillo motif-containing protein (SARM1) that inhibits TLR signalling and inflammasome activation upon KP infection^8,34^, affect the intracellular lipid A. However, infection of *il10^-/-^*, *aim2^-/-^*, *asc^-/-^* and *sarm1^-/-^* macrophages did not affect the ability of Kp52145 to produce the hydroxylated and palmitoylated hexa-acylated and hepta-acylated lipid A species (Fig S7).

To explain how TLR4-TRAM-TRIF signalling affects the production of the intracellular lipid A, we reasoned that this pathway may provide a cue for KP to remodel its lipid A. Our previous evidence demonstrating the importance of the acidic pH of the KCV led us to examine whether TLR4- TRAM-TIRF signalling controls the acidification of the KCV. To analyse the pH of the KCV, cells were loaded with LysoSensor after infection. LysoSensor serves as a ratiometric marker of vacuolar compartment pH (pKa of ∼4.2) as it is conjugated to the compound, 2-(4-pyridyl)-5-((4-(2- dimethylaminoethylaminocarbamoyl)-methoxy)-phenyl)oxazole that exhibits pH-dependent dual-excitation and dual-emission spectral peaks. Thus, LysoSensor produces a blue fluorescence in weakly acidic organelles and shifts to yellow in more acidic vacuoles. In wild-type macrophages, the KCV was acidic (Fig 6H) which was not the case in *tlr4^-/-^* and *tram-trif^-/-^* macrophages (Fig 6H). Quantification of the yellow fluorescence as a marker of KCV acidification showed that the fluorescence of the KCVs from *tlr4^-/-^* and *tram-trif^-/-^* was significantly lower than those of wild-type macrophages (Fig 6H), further substantiating that TLR4-TRAM-TRIF signalling is responsible for the acidification of the KCV.

Based on the connection between the pH of the KCV, the intracellular lipid A, and KP intracellular survival we hypothesized that absence of TLR4 should impair KP intracellular survival. Whereas no differences were observed in the adhesion of Kp52145 to wild-type and *tlr4^-/-^*(Fig 6I), the phagocytosis was lower in *tlr4^-/-^* cells than in wild-type macrophages (Fig 6J). Remarkably, the intracellular survival of Kp52145 was severely compromised in *tlr4^-/-^*macrophages (Fig 6K), providing experimental support to our hypothesis.

Altogether, this evidence supports a model in which TLR4-TRAM-TRIF signalling controls the KCV acidification, which is used by KP as a cue to produce a characteristic lipid A crucial for intracellular survival.

## DISCUSSION

Here, we have developed a method, we termed Intracell_lipidA_ method, to analyse the lipid A produced by intracellular KP which should be applicable to other intracellular Gram-negative bacteria. We demonstrate that KP remodels its lipid A within the KCV in mouse and human macrophages to survive intracellularly. KP lipid A is characterized by hexa and hepta-acylated species modified with palmitate and 2-hydroxylated. LpxL1, PagP and LpxO are the key enzymes governing the intracellular lipid A in a PhoPQ-dependent manner upon KP sensing of the acidic environment of the KCV. Remarkably, absence of TLR4 signalling impairs the production of the intracellular lipid A due to an increase in the KCV pH. KP intracellular survival is impaired in the absence of TLR4 signalling. The fact that TLR4 signalling also controls host defence responses against KP^29^ illustrates how a pathogen has evolved to require innate immune system cues for virulence.

The pathogen-containing vacuoles are harsh environments particularly those with an acidic pH. Research in this area mostly focuses on understanding how the pathogens deviate their vacuoles from the canonical maturation of the phagosome. A limited number of studies have ascertained the pathogens’ adaptations to this hostile environment although the evidence suggests that this environment has a profound impact upon the transcriptome^4–6^. Here, we demonstrate that intracellular KP remodels its lipid A to maintain membrane integrity in the KCV, illustrating an overlooked aspect when researching the intracellular lifestyle of Gram-negative bacterial pathogens.

In this work we have only focused on the lipid A given its relevance on bacterial physiology. However, it is tempting to hypothesize that KP may also remodel other surface structures, chiefly the peptidoglycan, to adapt to the environment within the KCV. Studies are ongoing to validate this hypothesis.

Our work determining the lipid A structure produced by intracellular bacteria (this work) and extracellular bacteria in vivo^13^ highlights the tremendous plasticity of the molecule and the relevance of these changes in the infection biology of KP. Although these structure-function studies have not been carried out for most Gram-negative pathogens and commensals there is enough evidence indicating that lipid A remodelling in the context of a mammalian system is a feature observed in other Gram-negative bacteria^35^. This evidence further emphasizes the limited value of probing LPS produced by bacteria grown in vitro to investigate in vivo host responses to Gram- negative bacterial infections. Furthermore, it is apparent that LPS from a single bacterium, for example *E. coli*, does not recapitulate the diversity and plasticity of the lipid A produced by pathogens in vivo and, therefore, caution must be exercised to generalize conclusions obtained using *E. coli* LPS.

Our results demonstrate that LpxO and PagP control the modifications of the intracellular lipid A; we have shown that they are also essential to remodel the lipid A produced by extracellular KP in the lungs of infected mice^13^. Not surprisingly, KP *lpxO* and *pagP* mutants are attenuated in vivo^10,13^, highlighting the importance of lipid A remodelling in KP infection biology. The fact that lipid A hydroxylation and palmitoylation also contribute to the virulence of other Gram-negative bacteria^36–38^ suggests that these lipid A modifications are the result of the host-pathogen co-evolution.

However, KP intracellular lipid A is distinct from those produced by other pathogens. The lipid A produced by *Shigella flexneri* residing in the cytosol of HeLa cells is hypo-acylated and contains no modifications^39^. Hypo-acylated species are also characteristic of the lipid A produced by *Burkholderia cenocepaia* in macrophages^17^ whereas the lipid A produced by *Salmonella typhimurium* within the *Salmonella*-containing vacuole (SCV) in mouse macrophages is hexa and hepta-acylated and it is extensively derivatized with 2-hydroxymyristate, palmitate, phosphoethanolamine, and/or aminoarabinose^15^. The two latter modifications were not found in KP intracellular lipid A. On the one hand, the differences between the lipid A structures produced by these pathogens could be explained by the different niches occupied by them intracellularly, the cytosol or a pathogen-containing vacuole, and, on the other hand, by the different environment within the vacuole. The pH of the *B. cenocepacia*-containing vacuole is almost neutral pH^40^ while it is acidic in the case of the SCV^41,42^ and KCV^9^ (this work). Nonetheless, the acidic pH is necessary but not sufficient to induce KP intracellular lipid A in vitro. Furthermore, growing KP in conditions reported to represent the environment within the SCV^4^ did not trigger the production of the lipid A found in the KCV. Together, this evidence demonstrates that the SCV and the KCV are indeed different despite some similarities. Future studies are warranted to design a medium that recapitulates the signals encountered by KP within the KCV.

A novel finding of our work is that TLR4-TRAM-TRIF signalling is required by KP to produce the intracellular lipid A, illustrating another angle of the host-pathogen arms race in which a pathogen exploits innate immune signalling for virulence. This work together with others from our laboratory substantiates that KP has evolved to hijack signalling by innate receptors, namely TLR4, NOD1, STING and NLRX1, to promote infection^43–46^. This strategy is radically different to those exploited by other pathogens who deploy toxins and bacterial effectors to control host defences. Mechanistically, we have demonstrated that the acidification of the KCV, a signal that controls the production of the intracellular KP lipid A, is impaired in the absence of TLR4 signalling. The mechanism is likely similar to that reported for TLR-dependent lysosomal acidification in dendritic cells upon recruitment of the vacuolar ATPase^47^. Supporting this notion, the acidification of the KCV is inhibited by bafilomycin A1, a specific and potent vacuolar ATPase inhibitor^28^. The role of TLRs in the maturation of phagosomes remains controversial^48–51^. Our study was not designed to address this controversy; however, our data clearly establishes that TLR4 signalling is required for the acidification of the KCV and has profound effects on the fate of intracellular KP. The fact that KP also exploits TLR4 signalling to rewire macrophage polarization^8^ further emphasizes the crucial role of TLR4 on the KP-macrophage interface.

Why does KP use TLR4 signalling to dictate its interactions with macrophages? A challenging scenario faced by KP is the need to adapt to extracellular and intracellular environments through the course of an infection. Precise regulation of the expression of virulence factors including systems needed to withstand host antimicrobial effectors, membrane remodelling, and metabolism is essential for KP to flourish in the tissues. Failure to do so most certainly results in decreased fitness and clearance by the innate immune system. We then posit the notion that TLR4-TRAM-TRIF signalling provides KP with a reliable input to determine its presence within the macrophage KCV. Linking the regulation of virulence downstream the activation of innate receptors is an efficient way of controlling the deployment of anti-host strategies only when and where needed.

## MATERIAL and METHODS

### Bacterial strains and growth conditions

All strains and plasmids used in this study are listed in Table S1. Strains were grown on LB agar from frozen glycerol stocks at 37°C. Isolated colonies were used to inoculate LB broth at 37°C on an orbital shaker (180 rpm). When adjusting pH of the LB, PCN or M9 was necessary, 20 mM MOPS was used to buffer media to pH 7 and 20 mM MES was used to buffer media to pH 5.0, 5.5 and 4.8 and adjusted using a Jenway 3510 Advanced Bench pH Meter. When indicated, antibiotics were used at the following concentrations: carbenicillin (Carb), 50 µg/ml; kanamycin (Km), 25 µg/ml; chloramphenicol (Cm), 30 µg/ml; trimethropin (Tmp) 100 µg/ml.

### Growth curve analyses

Analysis of growth kinetics was performed by inoculation of 5 μl of overnight bacterial culture into 250 μl sterile growth media, and incubated at 37 °C with continuous shaking using a Bioscreen C Automated Microbial Growth Analyser. Optical density (OD_600_) was measured and recorded at 20- minute intervals for 24 hours on five independent occasions.

### Cell culture

Human monocytes THP-1 (ATCC TIB-202) and the reporter cell line THP1-Dual (Invivogen, catalogue number thpd-nfis) were grown in Roswell Park Memorial Institute (RPMI) 1640 Medium (Gibco 21875) supplemented with 10% heat-inactivated fetal calf serum (FCS), 10 mM HEPES (Sigma), 100 U/ml penicillin, and 0.1 mg/ml streptomycin (Gibco) at 37 °C in a humidified 5% CO_2_ incubator.

Immortalised BMDM (iBMDM) from wild-type, *tlr2*^-/-^, *tlr4*^-/-^, *tlr2*^-/-^*tlr4*^-/-^, *myd88*^-/-^, *Tram*^-/-^*Trif*^-/-^, *irf3*^-/-^ mice on a C57BL/6 background, were obtained from BEI Resources (NIAID, NIH) (repository numbers NR-9456, NR-9457, NR-9458, NR-19975, NR-15633, NR-9568 and NR-15635, respectively). *IL-10*^-/-^, *aim2*^-/-^, *ifnar1*^-/-^, *sarm1*^-/-^ and *asc*^-/-^ iBMDMs were described previously^34,36^. iBMDM cells were grown in Dulbecco’s Modified Eagle Medium (DMEM; Gibco 41965) supplemented with 10% heat-inactivated FCS, 100 U/ml penicillin, and 0.1 mg/ml streptomycin (Gibco) at 37 °C in a humidified 5% CO_2_ incubator. Cells were routinely tested for *Mycoplasma* contamination.

### Infection conditions

Bacteria were grown in 5 mL LB at 37 °C, harvested at exponential phase (2500 x *g*, 20 min) and adjusted to OD_600_ of 1.0 in PBS (5×10^8^ CFU/ml). Infections were performed using a multiplicity of infection (MOI) of 100 bacteria per cell. Infections were performed in the same media used to maintain the cell-line without antibiotics and incubated at 37 °C in a humidified 5% CO_2_ incubator. THP-1 monocytes were differentiated into macrophages using phorbol 12-myristate 13-acetate (PMA) at 5 ng ml-1 at the time of seeding (2x10^5^ cells/ml) and infections were performed two days later. Infections of iBMDMs were performed the day after seeding (5x10^5^ cells/ml). To synchronise infections, plates were centrifuged at 200 x *g* for 5 min. In all macrophage infections extending beyond 1 h, the medium was replaced after 1 h with fresh medium containing 100 µg/ml gentamicin to kill extracellular bacteria. The extraction of lipid A from intracellular bacteria was done 2.5 h after the time of contact.

### Lipid A extraction using Ammonium Hydroxide/Isobutyric Acid

*In vitro* lipid A from *E. coli* and *K. pneumoniae* was extracted using the microextraction method^52^ and subjected to negative-ion matrix assisted laser desorption/ionisation time-of-flight (MALDI-TOF) mass spectrometric analysis. Briefly, 10 ml of fresh bacteria grown to mid exponential phase were washed twice with equal volumes of PB buffer to eliminate salt, LB, and other debris from the samples. The extraction was conducted on 10 mg of lyophilised bacteria. LPS was initially extracted by performing one or two washes using fresh 400 μl from a single-phase mixture of chloroform-methanol (1:2, v/v), followed by centrifugation at 2,000 x*g* for 15 minutes to remove most of the membrane phospholipids. Following centrifugation, the supernatant was discarded, and the pellet was washed a second time with 400 μl of chloroform-methanol-water. (3:2:0.25, v/v/v) and centrifuged again. To the dried cells, 400 μl of isobutyric acid-1 M NHCOH mixture (5:3, v/v) was added, followed by vortexing the samples. The crude LPS samples were subsequently hydrolysed at 95-100 °C for 2 hours, with rigorous vortexing every 15-20 minutes to cleave the Kdo linkage. After 2 hours, samples were placed on ice for 10-15 minutes to cease the hydrolysis reaction. Lipid A was harvested by centrifugation at 2,000 x*g* for 15 minutes, and the supernatant was diluted 1:1 with deionised water into fresh 2 ml screw-capped tubes, and frozen at -80oC, followed by lyophilisation.

### Bligh-Dyer lipid A extraction from intracellular bacteria

To extract lipid A from intracellular KP a modified Bligh-Dyer method was performed from a lyophilised or fresh bacterial suspension of KP, or intracellular infection^15^. 20x10^6^ cells were collected and the pellet was resuspended in 800 μl of PBS, followed by 2 ml of methanol and 1 ml of chloroform, and vortexed. The resulting mixture was incubated for 30 minutes at room temperature with occasional vortexing to allow the soluble phospholipids to separate from the insoluble LPS, DNA, and protein. The insoluble fraction was centrifuged at 2,000 x*g* for 15 minutes at room temperature. The pellet was washed twice more with a Bligh-Dyer suspension buffer in 4 ml of chloroform-methanol-water (1:2:0.8, v/v/v) with centrifugation at 2,000 x*g* for 15 minutes at room temperature. 3 ml of the chloroform-methanol-water supernatant was discarded, and the remaining 1 ml was used to suspend the pellet and subsequently transferred to a fresh 2 ml screw-capped tube and centrifuged 2,500 x *g* for 5 minutes at room temperature. Following centrifugation, the supernatant was discarded and resuspended in 400 μl of fresh isobutyric acid-1 M ammonium hydroxide continuing as described using Ammonium Hydroxide/Isobutyric Acid.

### Intracell_lipid A_ method to extract lipid A from intracellular bacteria

Mammalian cells were infected as described and upon collection were washed three times in sterile PBS and lysed with 0.05% saponin in PBS (w/v) at 37 °C for 5 minutes. Cells were washed in PB buffer 5 times and lyophilised in 500 μl of PB. Fresh or lyophilised KP or mammalian cells were suspended in fresh 200 μl of TRIzol Reagent and vortexed vigorously and incubated at room temperature for 15 minutes to homogenise the cell suspension. Following incubation, 20 μl of chloroform was added to encourage phase separation. The resulting mixture was vortexed vigorously for at least 10 seconds and incubated at room temperature for 10 minutes. The sample was then centrifuged at 12,000 x *g* for 10 minutes, separating the aqueous (containing the crude LPS) and organic phases. To harvest additional LPS molecules, 100 μl of deionised water was added to the organic phase and incubated at room temperature for 10 minutes, followed by centrifugation at 12,000 x *g* for 10 minutes. The aqueous phases were briefly vortexed, frozen at - 80°C, and lyophilised. Lyophilised crude LPS samples were briefly centrifuged at 12,000 x *g* and dissolved in 500 μl of mild acid hydrolysis buffer (1% SDS in 10 mM sodium acetate, pH 4.5), vortexed vigorously and heated at 100oC for 1 hour, with routine vigorous vortexing every 15 minutes. The mixture was immediately frozen at -80oC and lyophilised. To ensure SDS detergent contamination was at a minimum when analysed by the MALDI-TOF, the lyophilised samples were centrifuged briefly at 12,000 x *g* for 30 seconds to collect the lipid A. Firstly, 100 μl of deionised water was added to the lipid A, followed by a brief vortex, and 500 μl of acidified ethanol (100 μl 4 M HCl with 20 ml 95% ethanol) was also added. Secondly, the lipid A was vortexed again, and harvested by centrifugation at 2,000 x *g* for 15 minutes. Thirdly, the sample was then washed twice more with 500 μl of non-acidified 95% ethanol and centrifuged as before. Finally, the sample was suspended in 500 μl of PB buffer to ensure efficient freezing prior to freeze-drying, and the sample was lyophilised to yield purified lipid A ready for MALDI-TOF.

### MALDI-TOF analysis

A small volume of the solubilised lipid A was transferred to 1.5 ml microcentrifuge tubes and desalted with a few grains of ion-exchange resin (H+; Dowex 50W-X8) and briefly centrifuged. A 1 µl aliquot was deposited on a polished steel target plate for the dried-droplet method. An equal volume of 2,5-dihydroxybenzoic acid (2,5-DHB) matrix (Bruker Daltonics Inc.) saturated in 100 mM of citric acid Sigma Aldrich) or acetonitrile-0.1% trifluoroacetic acid (1:2 [vol/vol]) and allowed to air dry. Lipid A structural spectra were generated with a Bruker Autoflex Speed TOF/TOF mass spectrometer (Bruker Daltons Inc.) in negative reflectron mode with delayed extraction. All spectra were achieved with ion-accelerating voltage set at 20 kV and resulting spectra were generated with an average of 300 shots. A peptide calibration standard (Bruker Daltonics Inc.) was used to calibrate the MALDI-TOF mass spectrometer prior to analysis of each sample. Lyophilised *E. coli* MG1655 lipid A grown in LB broth at 37°C with identical extraction was used as an internal calibrant. Spectra are representative of at least three independent lipid A extractions.

### Tn7-based construction of *E. coli* strain expressing KP lipid A enzymes

To express Kp5245 *lpxO* in *E. coli*, the suicide vector pGPCTn7CCm_KpnLpxOCom was mobilized by conjugation into BN1-Δ*lpxL* harbouring pSTNSK-Tp plasmid. This plasmid encodes the transposase *tnsABCD* necessary for Tn7 transposition and it was introduced into *E. coli* by electroporation. Colonies were checked for resistance to Cm and sensitivity to Carb. As the ampicillin resistance cassette is located outside of the Tn7 region of the vector, Carb sensitivity denotes the integration of the Tn7 derivative at the *att*Tn7 site instead of incorporation of the vector into the chromosome. Confirmation of integration of the Tn7 transposon at the established *att*Tn7 site located 449 downstream of the *glmS* gene was verified by PCR using primers described in Table S2. pSTNSK-Tp from the recipient strains was cured by growing bacteria at 37 °C due to the plasmid thermosensitive origin of replication pSC101. Plasmid removal was confirmed by susceptibility to Tmp.

### Adhesion, phagocytosis and intracellular survival

iBMDMs were seeded in 12-well plates approximately 16 h before infection. Infections were performed at an MOI of 70 bacteria per cell. To enumerate the number of bacteria adhered to macrophages, after 1 hour of contact cells were washed twice with PBS, and lysed in 300 μl of 0.1% (w/v) saponin in PBS for 5 min at 37°C. Serial dilutions were plated in LB and the following day bacterial CFUs were counted. Results are expressed as CFU per ml. To determine the number of bacteria phagocytosed by the cells, after 1 hour of contact, cells were washed once with PBS and fresh medium containing gentamycin (100 μg/ml) was added to the wells. After 30 min, cells were washed three times with PBS, and lysed with saponin. Samples were serially diluted in PBS and plated in LB. After 24 h incubation at 37 °C, CFUs were counted and results expressed as CFUs per ml. To assess intracellular survival, 4.5 h after the addition of gentamycin, cells were washed three times with PBS and lysed with saponin. Serial dilutions were plated on LB to quantify the number of intracellular bacteria. Results are expressed as % of survival (CFUs at 5.5 h versus 1.5 h normalized to the results obtained in wild-type macrophages infected with Kp52145 set to 100%). All experiments were carried out with triplicate samples on at least five independent occasions.

### Assessment of bacteria membrane barrier

iBMDMs were seeded in 12-well plates (5x10^5^ cells/well) and infections were performed using a MOI of 70 bacteria per cell as described previously. The medium was replaced after 1 h with fresh medium containing 100 µg/ml gentamicin to kill extracellular bacteria and at 2.5 hours of infection cells were lysed in 300 μl of 0.1% (w/v) saponin in PBS for 5 min at 37°C. 100 µl of this bacterial suspension was then incubated in 96 well plates in the presence of PBS as vehicle control, 1% (w/v) NaDOC and/or 50 mM EDTA (pH 7.0) for 1 h at 37 °C. Samples were then serially diluted in PBS and plated in LB. After 24 h incubation at 37 °C, CFUs were counted and results expressed as CFUs per ml. Experiments were carried out with duplicate samples on three independent occasions.

### RNA isolation and RT-qPCR

Infections were performed in 6-well plates. Cells were washed two times with pre-warmed sterile PBS, and total RNA was extracted from the cells in 1 mL of TRIzol reagent (Ambion) according to the manufacturer’s instructions. The extracted RNA was further purified using MICOBEEnrich kit (Fisher Scientific) following the manufacturer’s instructions. Bacterial RNA was precipitated with sodium acetate and ethanol. RNA was quantified using a Nanovue Plus spectrophotometer (GE Healthcare Life Sciences). cDNA was generated by retrotranscription of 1 µg of total RNA using M-MLV reverse transcriptase (Invitrogen) and random primers (Invitrogen). Ten nanograms of cDNA were used as a template in a 5 µl reaction mixture from a KAPA SYBR FAST qPCR kit (Kapa Biosystems). Primers used are listed in Table S2. RT-qPCR was performed using a Rotor-Gene Q (Qiagen) with the following thermocycling conditions: 95 °C for 3 min for hot-start polymerase activation, followed by 40 cycles of 95 °C for 5 s and 60 °C for 20 s. Fluorescence of SYBR green dye was measured at 510 nm. Relative quantities of mRNAs were obtained using the ΔΔC_T_ method by using *rpoD* for gene normalization. RT-qPCR analyses were repeated on three independent occasions from three separate RNA extractions

### THP1 Dual

Reporter THP1-Dual cells (InvivoGen, catalogue thpd-nfis) were cultured Roswell Park Memorial Institute (RPMI) 1640 Medium (Gibco 21875) supplemented with 10% heat-inactivated fetal calf serum (FCS), 10 mM HEPES (Sigma), 100 U/ml penicillin, and 0.1 mg/ml streptomycin (Gibco) at 37 °C in a humidified 5% CO_2_ incubator. To maintain selection pressure, 10 μg/ml of Blasticidin and 100 μg/ml of Zeocin were added to the growth medium every other passage. THP-1 were seeded in 96 well plates using PMA at 20 ng/ml at the time of seeding (1x10^5^ cells/well) and media was replaced after 24h with RPMI-1640 without antibiotics. Infections were performed two days after seeding by adding 1.25 µl of a PBS bacterial suspension (MOI of 50:1). After incubation for 30 minutes at 37°C and 5% CO_2_, media was replaced by media containing gentamicin (100µg/ml). Cells were incubated for 16 hours before the culture supernatants were harvested and subjected to analysis of activity of a SEAP (NF-κB-activated expression) and luciferase activity (IFNAR signalling). All experiments were performed in duplicate, and three independent experiments were conducted.

### Quantification of cytokines

Infections were performed in 12-well plates (5 x 10^5^ cells per well) at an MOI of 100:1. After 5 h infection, supernatants were collected and spun down at 12000 x g for 2 min to remove any debris. TNF-α (#900-K54) and IP-10 (CXCL10) (#250-16) in the supernatants were quantified using ABTS ELISA Development Kits (PeproTech) according to the manufacturer’s instructions. All experiments were performed in duplicate, and three independent experiments were conducted.

### Assessment of the colocalization of the KCV with cellular markers

The protocol was adapted from Cano et al^9^. Briefly, wild-type iBMDMs (2x10^4^ per well) were grown on 13 mm circular coverslips in 24-well plates and were infected with Kp52145 and isogenic mutants harbouring pFPV25.1Cm^53^. After 1 hour of contact the coverslips were washed with PBS and gentamycin (100 μg/ml in DMEM medium) was added to kill extracellular bacteria. Cells were fixed at the indicated time points in the figure legends using 4% (w/v) paraformaldehyde in PBS pHC7.4 for 20 min at room temperature.

Cresyl violet acetate salt was used to label lysosomes^23^. Cresyl violet in fresh medium (5μM) was added to the cells 15 min before fixing the cells. The residual fluid marker was removed by washing the cells three times with PBS, followed by fixation. To determine the percentage of bacteria that co-localized with cresyl violet, bacteria located inside a minimum of 50 infected cells were analysed in each experiment.

LysoTracker Red DND-99 (Invitrogen) was used to label acidic organelles following the instructions of the manufacturer. 0.5µM Lysotracker Red DN99 was added to the tissue culture medium 30 min before fixing the cells. The residual fluid marker was removed by washing the cells three times with PBS, followed by fixation.

For Rab14 staining, coverslips were washed with PBS and permeabilized with 0.1% (w/v) saponin (Sigma) in PBS for 30 min. Coverslips were then incubated for 120 min with anti-Rab14 (4 μg/ml in 0.1% (v/v) horse serum (Gibco), 0.1% (w/v) saponin in PBS; clone D-5, murine IgG1, sc- 271401, Santa Cruz), washed with PBS, followed by a 45 min incubation with anti-mouse IgG H&L labelled with AlexaFluor 647 (10 μg/ml in 0.1% (v/v) horse serum (Gibco), 0.1% (w/v) saponin in PBS, polyclonal, donkey IgG, ab150111, Abcam). Coverslips were washed with PBS, and then incubated with anti-Lamp1 (1 μg/ml in 0.1% (v/v) horse serum (Gibco), 0.1% (w/v) saponin in PBS, clone 1D4B, rat IgG2a, sc-19992, Santa Cruz) for 20 min, washed with PBS, and incubated for 20 min with anti-rat IgG H&L labelled with AlexaFluor 568 (10 μg/ml in 0.1% (v/v) horse serum (Gibco), 0.1% (w/v) saponin in PBS, polyclonal, goat IgG, A11077, Life Technologies). To determine the percentage of the Lamp1 positive KCV that co-localized with Rab14, KCVs of at least 100 infected cells from three independent experiments were analysed.

Coverslips were mounted with ProLong Gold antifade mountant (Invitrogen) and visualised on the Leica DMi8 Confocal microscope. Experiments were carried out in duplicate in three independent occasions.

### Immunoblotting

Macrophages were seeded in 6-well plates for 24 h before infection. Cell lysates were prepared in lysis buffer (1x SDS Sample Buffer, 62.5 mM Tris-HCl pH 6.8, 2% w/v SDS, 10% glycerol, 50 mM DTT, 0.01% w/v bromophenol blue). Proteins were resolved on 8, 10 or 12% SDS-PAGE gels and electroblotted onto nitrocellulose membranes. Membranes were blocked with 3% (wt/vol) bovine serum albumin in TBS-Tween (TBST), and specific antibodies were used to detect protein using chemiluminescence reagents and a G:BOX Chemi XRQ chemiluminescence imager (Syngene).

Phosphorylated AKT1/2/3 was detected using anti-phospho-AKT1/2/3 (Ser 473) (anti-rabbit, 1:1000; sc-33437, Santa Cruz). Immunoreactive bands were visualized by incubation with HRP-conjugated IgG Secondary antibody (goat anti-rabbit, 1:5000; #170-6515, Bio Rad). To ensure that equal amounts of proteins were loaded, blots were re-probed with α-tubulin (1:3000; #T9026, Sigma- Aldrich). To detect multiple proteins, membranes were re-probed after stripping of previously used antibodies using a pH 2.2 glycine-HCl/SDS buffer.

### Assessment of KCV acidification

5x10^5^ iBMDMs were plated in a 10 mm Glass Bottom Culture 35 mm petri dish (MATEK corporation, P35G-0-14-C) and incubated 12–16 h at 37 °C and 5% CO_2_. The next day, cells were infected with GFP tagged KP as described above. 15 minutes prior to the indicated infection time points, cells were washed twice with PBS and incubated in DMEM with LysoSensor Green DND- 189 (1μM; Molecular Probes, Invitrogen, L7535). The cells were then rinsed twice in PBS and were immediately analyzed by confocal microscopy. During image acquisition, the cells were maintained at 37 °C and 5% CO_2_ in a temperature-controlled CO_2_ chamber on the microscope. To determine if the vacuole was neutral or acidic, KCVs of at least 100 infected cells from five independent experiments were analysed.

### Statistical analysis

Statistical analyses were performed using one-way analysis of variance (ANOVA) with Bonferroni corrections, *P* values of <0.05 were considered statistically significant. Normality and equal variance assumptions were tested with the Kolmogorov-Smirnov test and the Brown-Forsythe test, respectively. All analyses were performed using GraphPad Prism for Windows (version 10.4.2) software.

## ACKNOWLEDGEMENTS

We thank the members of the J.A.B. laboratory for their thoughtful discussions and support with this project. T.L.B. is the recipient of a Ph.D. fellowship funded by the Department for Employment and Learning (Northern Ireland, UK). This work was supported by Biotechnology and Biological Sciences Research Council (BBSRC, BB/P006078/1 and BB/P020194/1) and Medical Research Council (MRC, MR/V032496/1) funds to J.A.B.

## AUTHOR CONTRIBUTIONS

Conceptualization, J.A.B., T.L.B. and J. sP; Investigation, T.L.B., J. sP., R.L., N.Z., K. W-O’F. Resources, S.J.H., G.M., and A.K.; Funding acquisition, J.A.B.; Writing original draft, J.A.B., J. sP., T.L.B. ; Writing-Review and Editing J.A.B., J. sP., T.L.B and A.K. Supervision, J.A.B and J. sP.

## DATA AVAILABILITY

All relevant data are presented in the manuscript, the supplementary information and the respective source files. Should any raw data files be needed in another format, they are available from the corresponding author.

## SUPPLEMENTARY FIGURES

**Figure S1.** Lipid A structures produced by *K. pneumoniae*. The mass-to-charge ratios (*m/z*) of each lipid A species is shown as well as the acyl chains esterifying the 2′ and 3′ R-3-hydroxymyristoyl groups. The lipid A modifications with palmitate (C_16_), and 4CaminoC4CdeoxyCLCarabinose (L-Ara4N) and phosphoethanolamine (pEtN) to the 4′ and 1-phosphate, respectively, are also depicted.

**Figure S2.** Glycolipids extraction from non-infected macrophages by different methods. A. Negative-ion MALDI-TOF mass spectrometry spectra of glycolipid purified from non-infected iBMDMs by the Bligh-Dyer method. B. Negative-ion MALDI-TOF mass spectrometry spectra of glycolipid purified from non-infected iBMDMs by the Tri Reagent method. C. Negative-ion MALDI-TOF mass spectrometry spectra from non-infected iBMDMs by the Intracell_lipid A_ method developed in this study. D. Negative-ion MALDI-TOF mass spectrometry spectra from Kp52145 grown in LB by the Intracell_lipid A_ method. E. Negative-ion MALDI-TOF mass spectrometry spectra of lipid A purified from intracellular KP35. In all panels data represent the mass-to-charge ratios (*m/z*) of each lipid A species detected and are representative of three extractions.

**Figure S3.** Proposed structures of the lipid A produced by intracellular *K. pneumoniae*. The mass-to-charge ratios (*m/z*) of each lipid A species is shown as well as the acyl chains esterifying the 2′ and 3′ R-3-hydroxymyristoyl groups. The lipid A modification with palmitate (C_16_) is also depicted.

**Figure S4.** Characterization of the enzymatic activity of LpxL1. A. Negative-ion MALDI-TOF mass spectrometry spectra of lipid A purified from *E. coli* BN1. B. Negative-ion MALDI-TOF mass spectrometry spectra of lipid A purified from *E. coli* BN1 *lpxL* mutant (strain BN1-Δ*lpxL*). C. Negative-ion MALDI-TOF mass spectrometry spectra of lipid A purified from *E. coli* BN1 *lpxL* mutant (strain BN1-Δ*lpxL*) encoding KP *lpxO* ntegrated into the *att* Tn7 site of teh chromosome. D. Negative-ion MALDI-TOF mass spectrometry spectra of lipid A purified from *E. coli* BN1 *lpxL* mutant (strain BN1-Δ*lpxL*) encoding KP *lpxO* ntegrated into the *att* Tn7 site of teh chromosome and complemented with KP LpxL1. C. Negative-ion MALDI-TOF mass spectrometry spectra of lipid A purified from *E. coli* BN1 *lpxL* mutant (strain BN1-Δ*lpxL*) and complemented with KP *lpxL1*. In all panels data represent the mass-to-charge ratios (*m/z*) of each lipid A species detected and are representative of three extractions.

**Figure S5.** Fitness of lipid A mutants. Growth kinetics of Kp52145 (black)), *lpxO* and *pagP* mutants (red and blue, respectively) cultured in: A. LB buffered pH 7.0 over 24 h at 37°C. B. LB buffered pH 5.5 over 24 h at 37°C. C. 2% glucose M9 minimal media supplemented with thiamine and MgSO4 (M9-glucose) buffered pH 7.0 over 24 h at 37°C. D. 2% glucose M9 minimal media supplemented with thiamine and MgSO4 (M9-glucose) buffered pH 5.5 over 24 h at 37°C. E. 2% glucose M9 minimal media supplemented with thiamine and MgSO4 (M9-glucose) buffered pH 4.8 over 24 h at 37°C. F. 2% glucose M9 minimal media supplemented with thiamine, MgSO4 (M9-glucose) and 10 µM MgCl_2_ and 10 buffered pH 5.0 over 24 h at 37°C. G. PCN medium containing 25 mM K_2_HPO_4_/KH_2_PO_4_ buffered to pH 5.0 over 24 h at 37°C. Values are presented as the mean ± SD of three independent experiments measured in triplicate.

**Figure S6.** Bafilomycin treatment abrogates the overlap of the KCK with LysoTracker. Immunofluorescence confocal microscopy of the coClocalisation of the KCV with LysoTracker in infected wild-type iBMDMs with Kp52145 harbouring pFPV25.1 and treaded with vehicle control (DMSO) or 100 nM befilomycin. A1. Bafilomycin was added after 1 h time of contact and maintained for the duration of the experiment. LysoTracker was added to cells 30 min before fixation. The images were taken 2.5 post gentamycin treatment to kill extracellular bacteria. Images are representative of duplicate coverslips of three independent experiments.

**Figure S7.** Lipid A produced by *K. pneumoniae* in macrophages deficient in innate proteins. Negative-ion MALDI-TOF mass spectrometry spectra of lipid A purified from: A. Intracellular Kp52145 in *il10^-/-^* iBMDMs. B. Intracellular Kp52145 in aim2*^-/-^* iBMDMs. C. Intracellular Kp52145 in *asc*^-*/-*^ iBMDMs. D. Intracellular Kp52145 in sarm1*^-/-^* iBMDMs. Data represent the mass-to-charge ratios (*m/z*) of each lipid A species detected and are representative of three extractions.

