## Supplementary figures and images for "*Klebsiella pneumoniae* remodels its Kdo_2_-lipid A in a TLR4-dependent manner to adapt to the macrophage intracellular environment"

### Figure S1

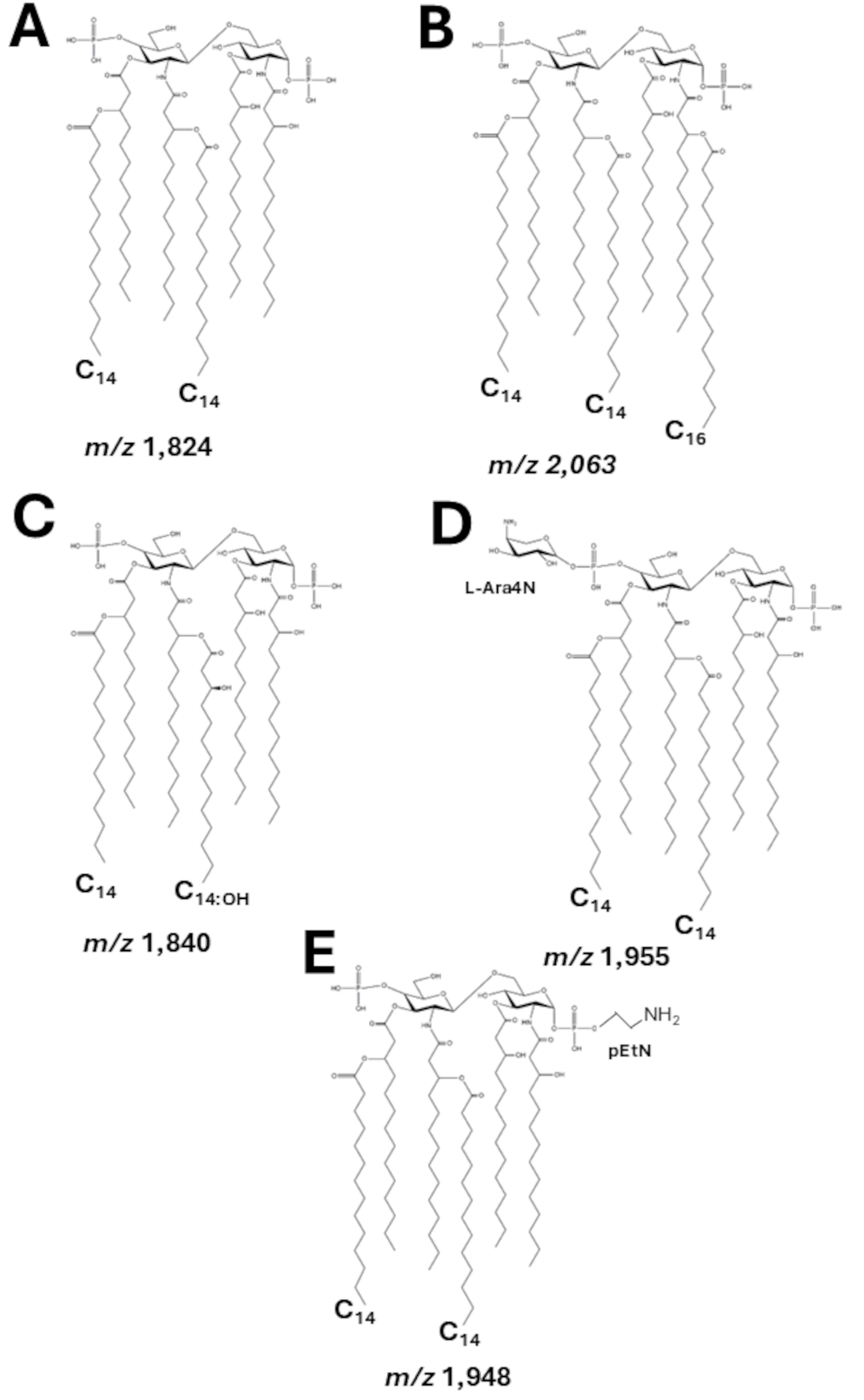

### Figure S2

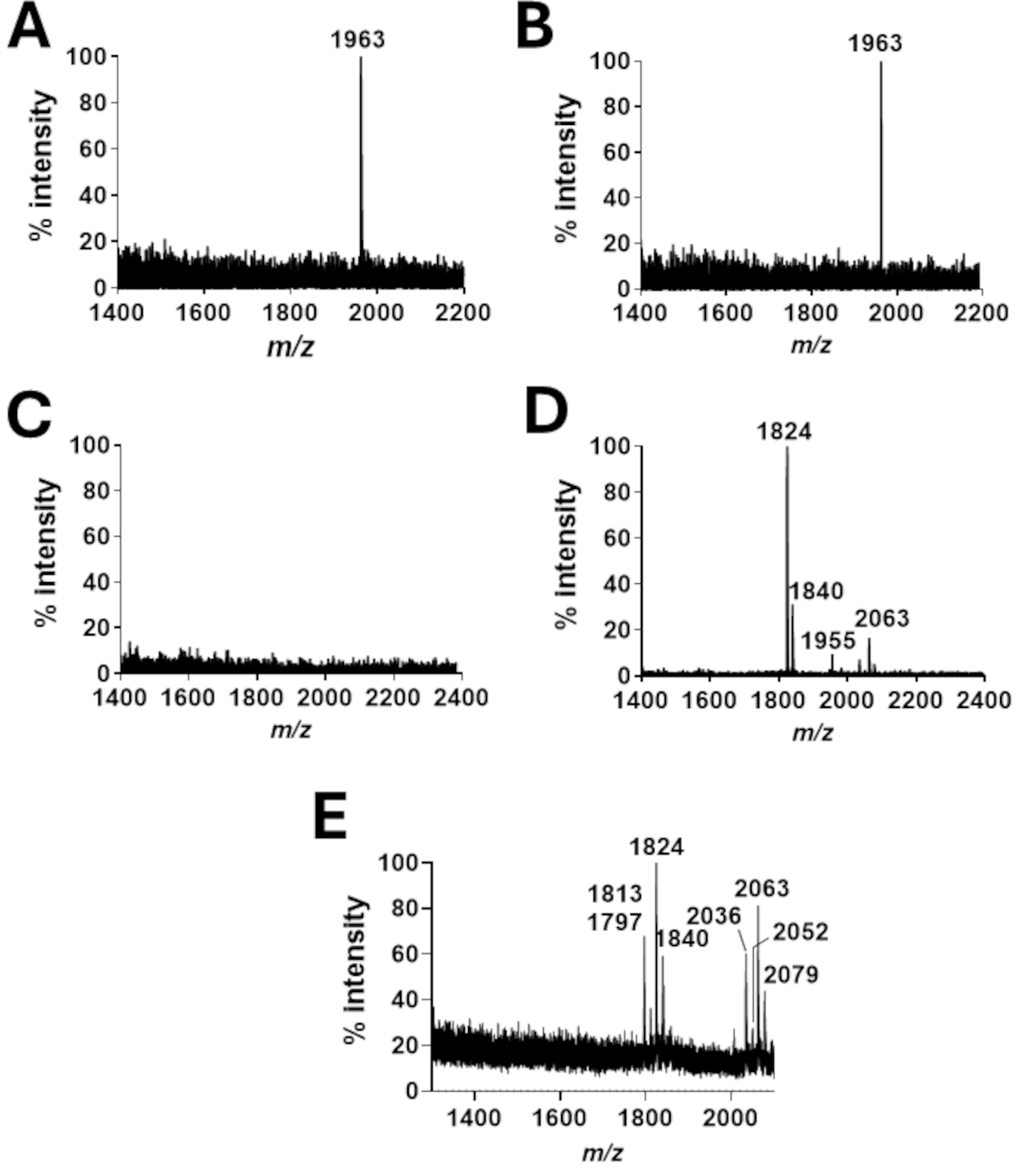

### Figure S3

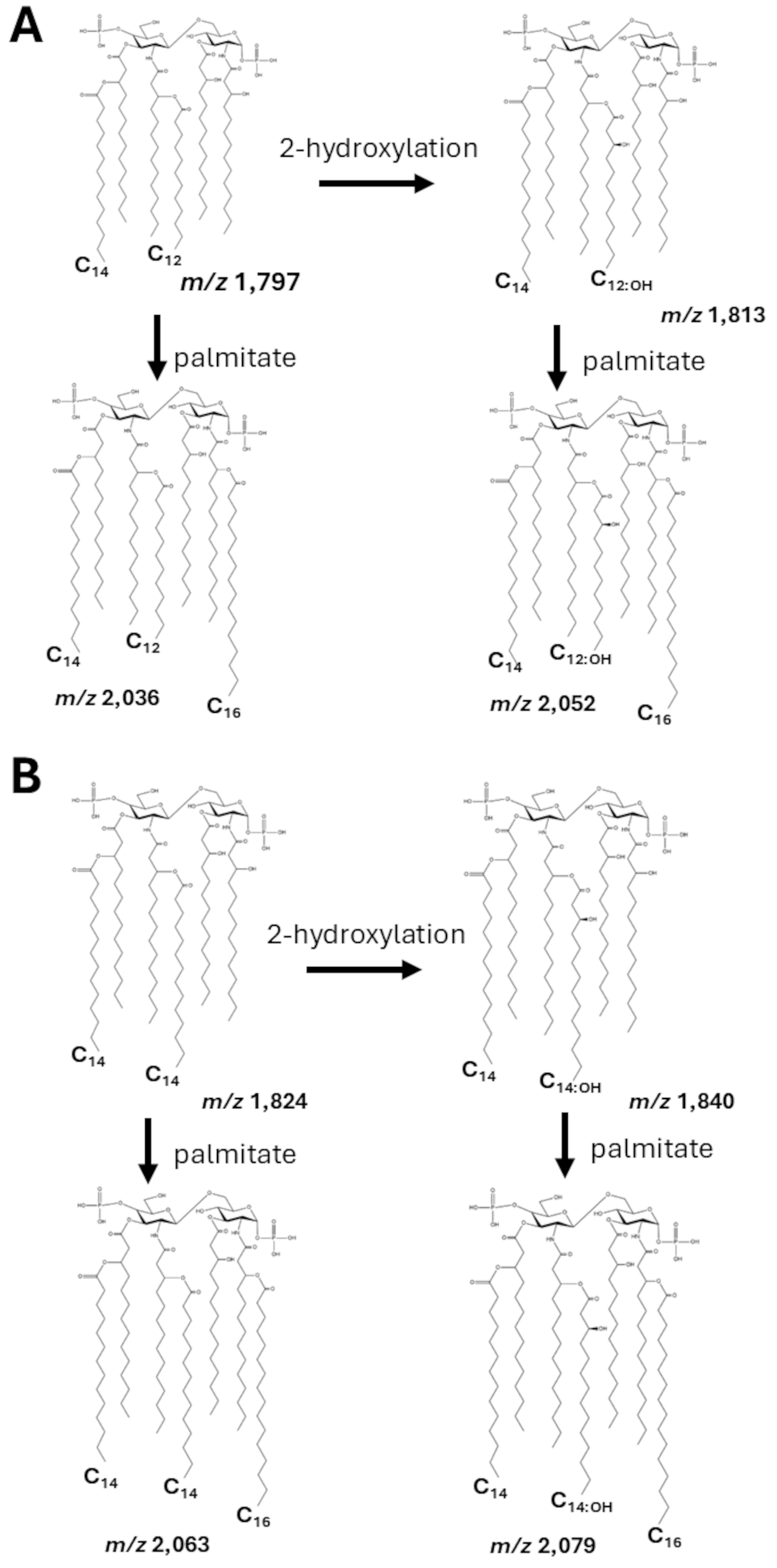

### Figure S4

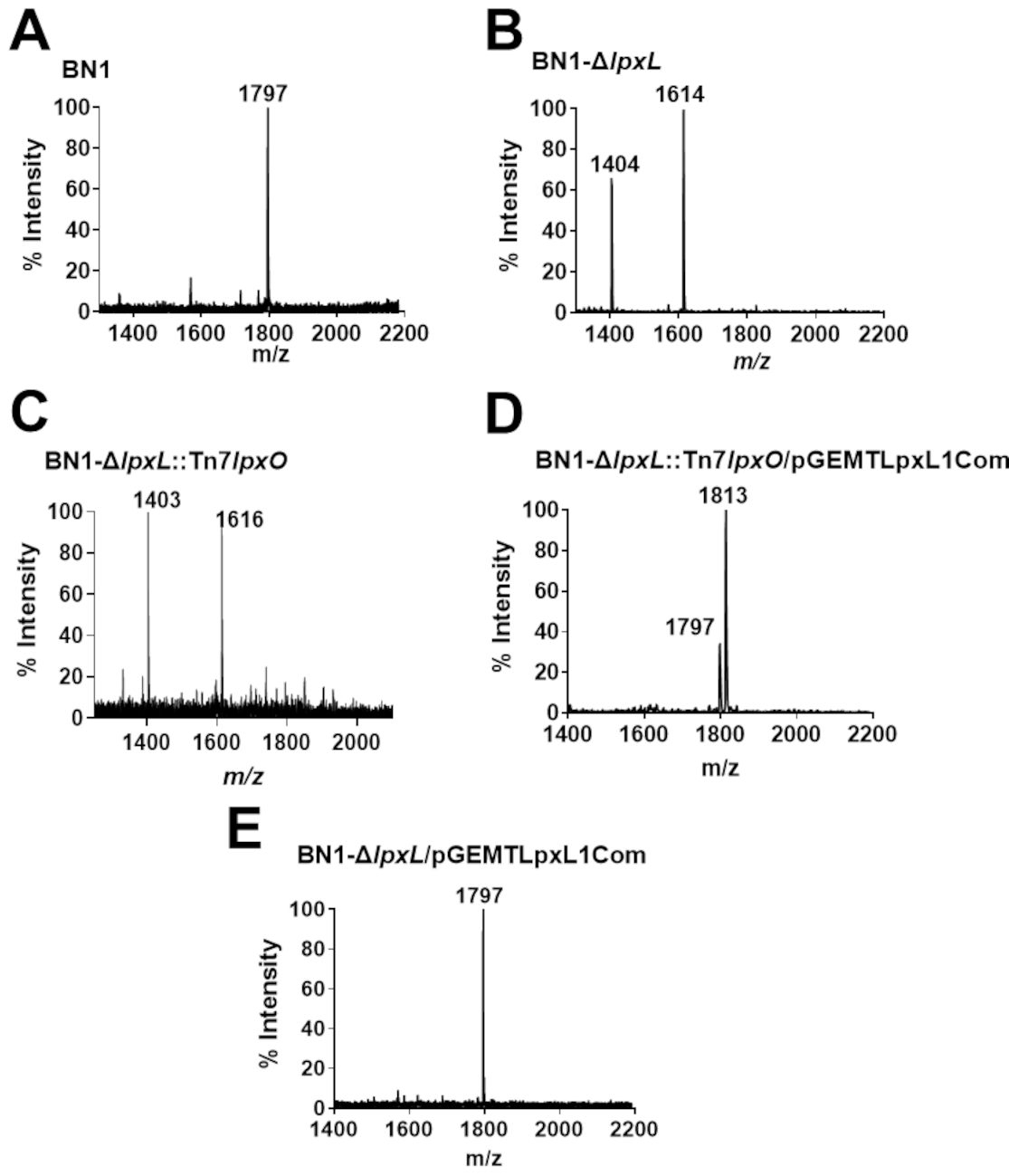

### Figure S5

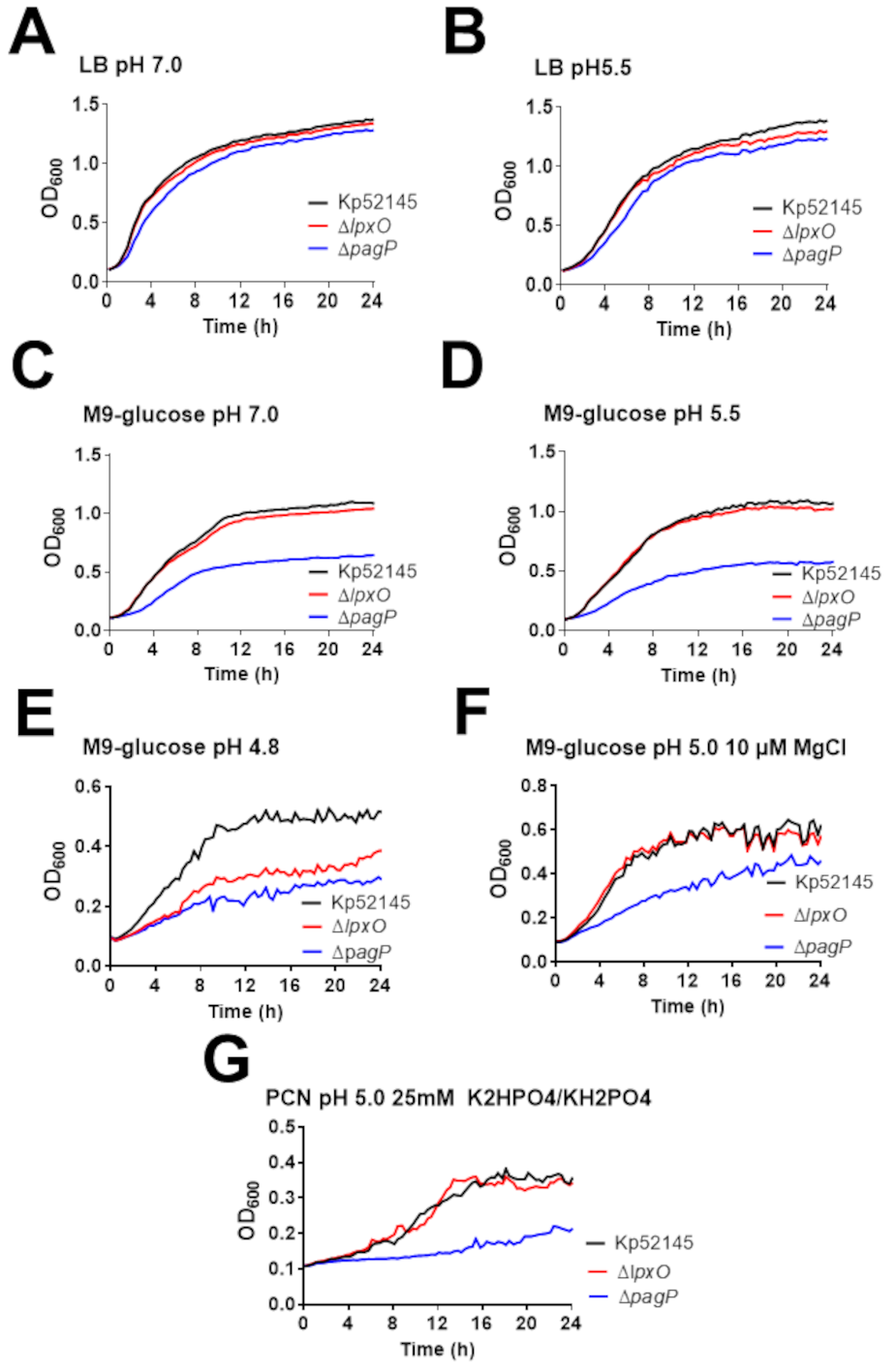

### Figure S6

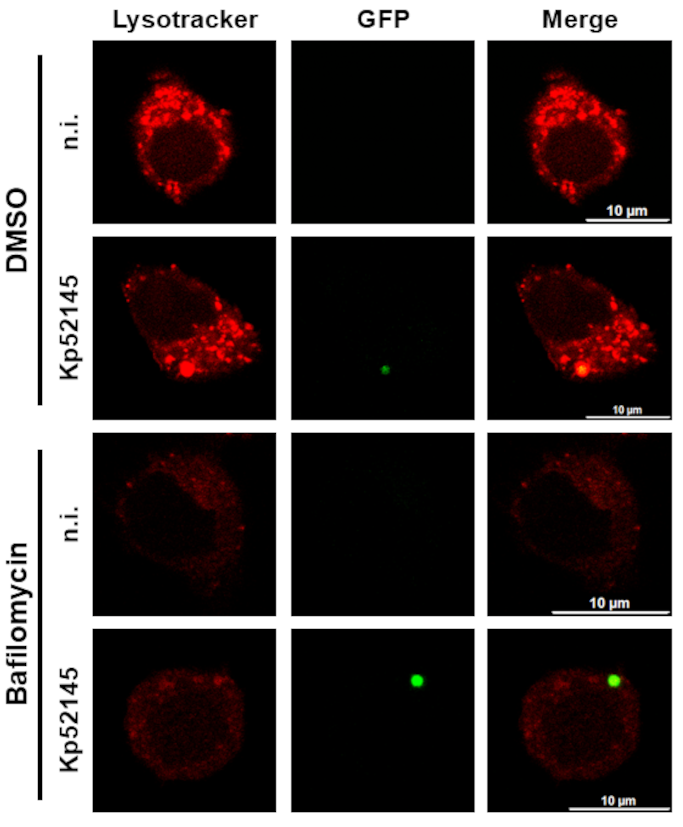

### Figure S7

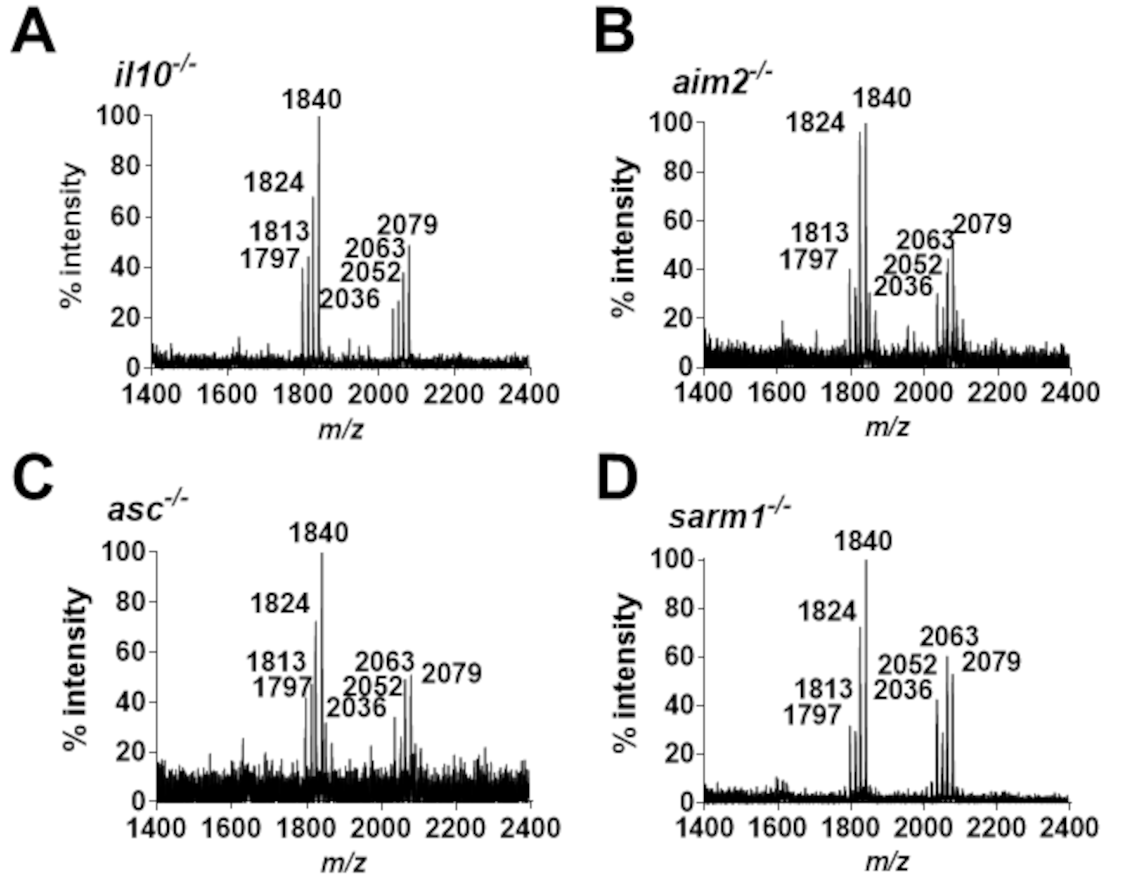
