## Supplementary material for "*Klebsiella pneumoniae* remodels its Kdo_2_-lipid A in a TLR4-dependent manner to adapt to the macrophage intracellular environment": Table S1

**Table S1. Strains and plasmids used in this study.**

| **Bacterial strain** | **Genotype or comments** | **Source** |
| --- | --- | --- |
| CIP 52.145 | Kp52145; Clinical isolate; serotype O1:K2;  sequence type ST66 | ^1^ |
| KP35 | Carbapenem resistant clinical isolate from a patient with bacteraemia sequence type ST258 | ^2^ |
| 52145-Δ*lpxL1* | Kp52145, ∆*lpxL1::FRT*; the *lpxL1*gene was inactivated | ^3^ |
| 52145-Δ*lpxL1*Com | Kp52.145 ΔlpxL1; Tn7Km_lpxL1 was  integrated into the attTn7 site, Kmr^R^ | ^3^ |
| 52145- Δ*lpxO* | Kp52145 with the *lpxO* gene inactivated | ^4^ |
| 52145- Δ*lpxO*Com | Kp52145, Δ*lpxO*; Tn7Cm_KpnlpxOCom  integrated into attTn7 site, Cm^R^ | ^4^ |
| 52145-Δ*pagP* | Kp52145 with the *pagP* gene inactivated | ^5^ |
| 52145-Δ*pagP*Com | Kp52145 Δ*pagP*; *pagP* inserted intp intergenic region *pgpA* and *yajO* KP genome (to be described). | This work |
| 52145- Δ*phoQ*GB | Kp52145 Δ*phoQ*::KM-GenBlock | ^6^ |
| 52145- Δ*phoQ*GBCom | Kp52145 Δ*phoQ*::KM-GenBlock complemented with pGP-Tn7-Cm_KpnPhoPQCom; *phoPQ* activity was restored | ^4^ |
| 52145- Δ*pmrAB* | Kp52145 with *pmrAB* genes inactivated | ^6^ |
| BN1 | *E. coli* W3110 Δ*eptA* Δ*lpxT* Δ*pagP* | ^7^ |
| BN1- Δ*lpxL* | BN1 Δ*lpxL*::FRT; the *lpxL* gene was  inactivated | ^3^ |
| BN1-Δ*lpxL::*Tn7lpxO | BN1Δ*lpxL*::FRT; Tn7-Cm_KpnLpxOCom  integrated into *att*Tn7 site; Cm^R^ | This work |
| BN1-Δ*lpxL::*Tn7lpxO/pGEMTlpxL1Com | BN1Δ*lpxL*::FRT; Tn7-Cm_KpnLpxOCom harbouring K52145 *lpxL1* cloned in to pGEM-T Easy | This work  ^3^ |
| **Plasmids** |  |  |
| pGEMTlpxL1Com | pGEM-T Easy containing Kp52145 Δ*lpxL1*; Amp^R^ | ^3^ |
| pGP-Tn7-Cm_KpnLpxOCom | To complement Kp52145 *lpxO* mutant by insertion into *att*Tn7 site of the genome, Amp^R^, Cm^R^ | ^4^ |
| pSTNSK-Tp | pSTNSK-Tp containing a transposase for Tn7 integration; Km^R^ , Tmp^R^ | ^8^ |
| pGPI-pgpA-pagP | Suicide vector, R6Kɣ origin of replication, Mob^+^ , carries a I-SceI endonuclease site; to integaret Kp52145 pagP into the intergenic region *pgpA* and *yajO,* Tmp^R^ | To be described |

Km^R^ , kanamycin resistant; Tmp^R^ , trimethoprim resistant; Amp^R^ , ampicillin resistant; Cm^R^ , chloramphenicol resistant.
