## Supplementary material for "*Klebsiella pneumoniae* remodels its Kdo_2_-lipid A in a TLR4-dependent manner to adapt to the macrophage intracellular environment": Table S2

**Table S2. Primers used in this study**

| **Method and target gene** | **Primer** | **Sequence (5’-3’)** |
| --- | --- | --- |
| Tn7 integration in *E. coli* |  |  |
|  | E.coli_glmS_UP | TCG ACT GGG CGT ACA AAA CC |
|  | E.coli_glmS_DWN | CGG GAA ACC ATA CCG GAG TT |
|  | E.coli_pTn7L_F1 | ATT AGC TTA CGA CGC TAC ACC C |
|  | E.coli_pTn7R_R1 | CAC AGC ATA ACT GGA CTG ATT TC |
| **qPCR** |  |  |
| *rpoD* | Kpn_RpoD_F1 | CCG GAA GAC AAA ATC CGT AA |
|  | Kpn_RpoD_R1 | CGG GTA ACG TCG AAC TGT TT |
| *lpxL1* | Kpn_LpxL1_F1 | TAT CGC CCC AAC GAC AAC C |
|  | Kpn_LpxL1_R1 | AAA CAG TGG GGC GAA GAC G |
| *lpxO* | Kpn_LpxO_F2 | TCG CTA CGC TTT CAT CTG GG |
|  | Kpn_LpxO_R2 | TCA GGC GAT TCT CAC CAC TG |
| *pagP* | Kpn_PagP_F2 | CGC TGG GAT GAG AAA GGG AA |
|  | Kpn_PagP_R2 | CGT GAC GCC GAG GGT ATA G |
